# Cryptic arsenic cycling controls oxygenic photosynthesis in Precambrian-analog microbial mats

**DOI:** 10.1101/2024.04.27.591451

**Authors:** Andrea Castillejos Sepúlveda, Daan R. Speth, Carolin Kerl, Daniel Doherty-Weason, Federico A. Vignale, Kinga Santha, Wiebke Mohr, Arjun Chennu, Elisa Merz, Dirk de Beer, Maria E. Farias, Judith M. Klatt

**Affiliations:** Microsensor Group, Max Planck Institute for Marine Microbiology, Bremen, Germany; Biogeochemistry Group, Max Planck Institute for Marine Microbiology, Bremen, Germany; Division of Microbial Ecology, Centre for Microbiology and Environmental Systems Science, University of Vienna, Vienna, Austria; Environmental Geochemistry, Bayreuth Center for Ecology and Environmental Research (BayCEER), University of Bayreuth, 95440 Bayreuth, Germany; Microcosm Earth Center, University of Marburg & Max Planck Institute for Terrestrial Microbiology, 35032 Marburg, Germany; Center for Synthetic Microbiology (SYNMIKRO), 35032 Marburg, Germany; Biogeochemistry Group, Department of Chemistry, University of Marburg, 35032 Marburg, Germany; European Molecular Biology Laboratory - Hamburg Unit, Hamburg 22607, Germany; Australian Institute of Marine Science, Townsville, Australia; University of Konstanz, Plant Ecophysiology, University of Konstanz, Germany; PUNABIO S.A. Campus USP-T Av. Solano Vera y Camino a Villa Nougués, San Pablo, Tucumán, Argentina

## Abstract

The delayed rise of atmospheric oxygen, despite the early evolution of oxygenic photosynthesis (OP), remains a central puzzle in Earth history. Numerous ecological and geochemical constraints on OP have been proposed, but the role of environmental stressors at the physiological and ecosystem level is poorly understood. Here we show that Chl-*f*-harboring cyanobacteria in a high-altitude Andean microbial mat – an analog for Precambrian ecosystems – switch from OP to arsenite-driven anoxygenic photosynthesis (AP) under high light. Using microsensor profiling, mat incubations, and metatranscriptomics, we show that this shift is triggered by the accumulation of reactive oxygen species (ROS), especially hydrogen peroxide, which suppresses OP. Instead of ceasing activity, cyanobacteria reroute electron flow, using arsenite as the electron donor to sustain photosynthesis while avoiding both intracellular ROS from OP and extracellular ROS from aerobic arsenite oxidation. This switch is reversible and coordinated with diel cycles of light and arsenic speciation, sustained by a cryptic arsenic redox cycle, continuously regenerating arsenite for AP. Although the enzymatic basis remains unresolved, these findings reveal a hidden layer of metabolic plasticity in cyanobacteria and suggest that oxidative stress-responsive metabolic shifts may have supported early phototroph survival while limiting oxygen release – potentially contributing to Earth’s protracted oxygenation.

## Main

The evolution of oxygenic photosynthesis (OP) in cyanobacteria was a transformative event in Earth’s history, fundamentally reshaping our planet. OP likely emerged during the Archean eon (3.7–2.5 billion years ago), within microbial mats – layered microbial communities that dominated shallow waters^1–3^. Before the rise of OP, light-driven microbial primary production relied on anoxygenic photosynthesis (AP), which uses reduced substances such as ferrous iron or sulfide as electron donors for carbon fixation^4,5^. These metabolisms shaped early biogeochemical cycles and may have delayed the proliferation of OP across Earth’s surface^6–8^.

Intriguingly, some modern cyanobacteria are capable of both OP and AP, using sulfide as preferred electron donor when available^9–14^. Thus, it has been proposed that even early cyanobacteria may have already transitioned between modes of photosynthesis in response to environmental cues. However, the evolutionary timeline of photosynthetic versatility and the extent to which such flexibility influenced the evolutionary trajectory of OP remains unclear^15,16^.

Arsenite (AsO_3_ ^3-^, As(III)), a reduced and bioavailable form of arsenic, was abundant in Archean and Proterozoic environments due to intense volcanic activity^17,18^. Like sulfide, As(III) can serve as an electron donor for AP in obligate anoxygenic phototrophs and can fuel a complete microbial arsenic redox cycle^19,20^. In cyanobacteria, arsenic and sulfur metabolism are co-regulated, suggesting an evolutionary connection^21^. Despite this link and arsenic’s environmental relevance, the role of arsenic in modulating early photosynthesis remains poorly understood.

One likely setting for the emergence of cyanobacteria is microbial mats, where they would have navigated steep chemical gradients, intense ultraviolet (UV) radiation, and reduced, often toxic, compounds like As(III). We hypothesize that As(III) availability may have influenced the balance between AP and OP – either by providing an alternative electron donor or by chemically interfering with the efficiency and evolution of OP. Arsenic could have thereby acted as both a metabolic substrate and a physiological stressor, shaping the evolutionary trajectory of early phototrophs.

To address this, we studied microbial mats from Laguna Pozo Bravo, a high-altitude hypersaline lake in the Central Andes that serves as a modern analogue to Precambrian mat ecosystems. Combining high-resolution microsensor profiling with incubation experiments and multi-omics approaches, we investigated how local arsenic speciation affects photosynthesis within these layered communities. Our findings provide new insights into the metabolic plasticity of cyanobacteria under extreme environmental stress and reveal previously unrecognized mechanisms by which arsenic may have constrained the early expansion of OP.

### A Precambrian-analog microbial ecosystem in the high Andes

Microbial mats are among the oldest known ecosystems on Earth, and modern analogs found in extreme environments offer valuable windows into early microbial life Earth^3,22–26^. Laguna Pozo Bravo, a hypersaline lake in the Central Andes (∼3300 m elevation), is one such system^27–29^. Its high UV radiation, extreme temperature fluctuations, and elevated arsenic levels – due to the volcanic origin of the surrounding geology – recreate conditions believed to be common in shallow aquatic environments of the Precambrian eon^27,29,30^.

In February 2019 and January 2023 the lagoon floor was extensively covered by cohesive, light pink microbial mats, thriving underneath a hypersaline (137.25 g L^-1^) water column with a stable temperature around 20°C. The mats formed stable sheets atop coarse sediment and displayed clear vertical stratification into four visible layers: a light pink upper crust rich in calcium carbonate, a green layer, a red layer, and a brown layer (Fig. 1a, inset).

**Figure 1.**
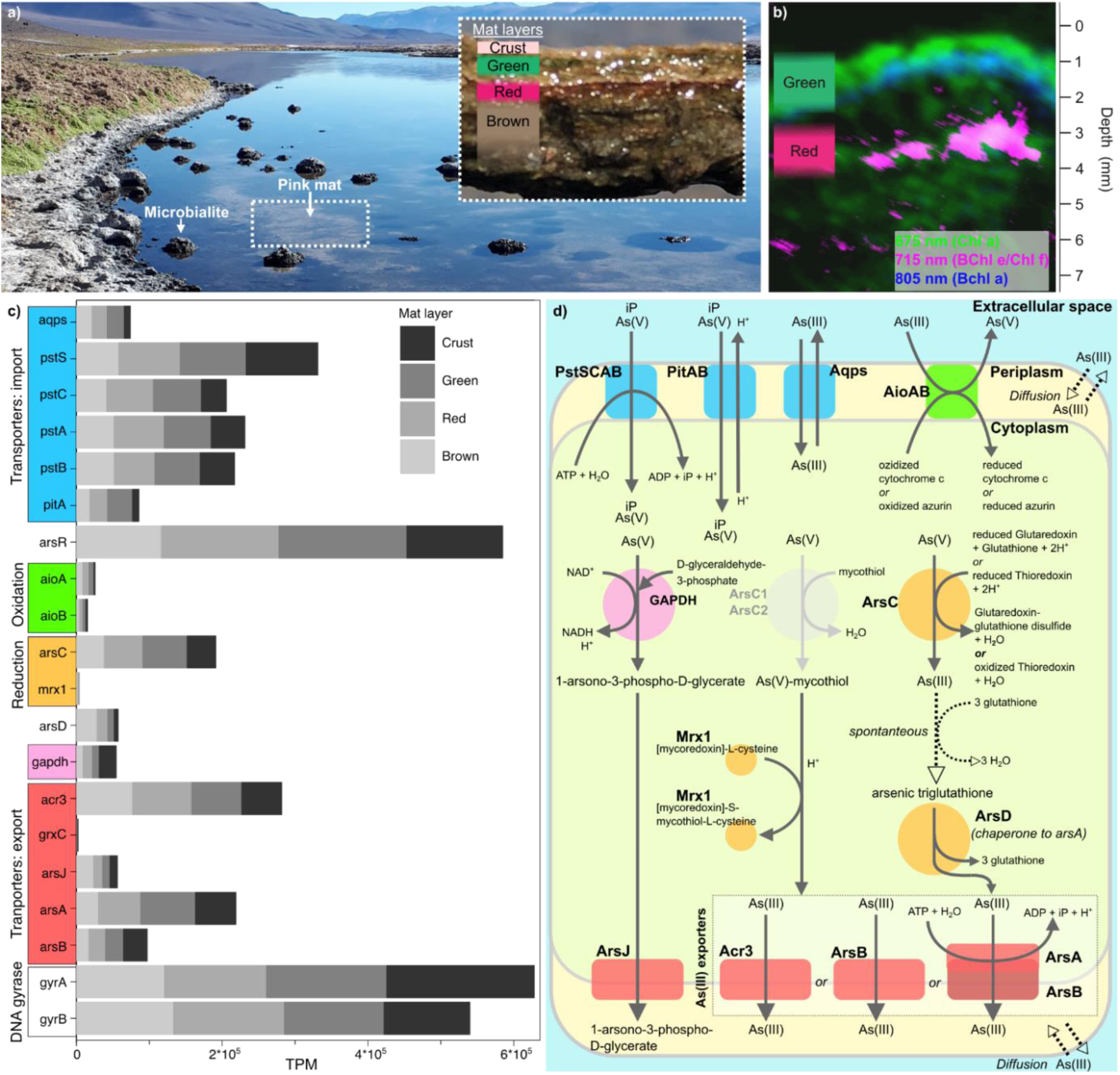
The microbial mats of Laguna Pozo Bravo. a) Microbialites and microbial mats (pink and lithifying gray) found in Pozo Bravo. Inset is a pink microbial mat cross section, with layers indicated. b) Distribution of selected pigments in the top 7 mm of Pozo Bravo microbial mat, captured by hyperspectral imaging, based on the second derivative of reflectance spectra around 675 nm, 715 nm, and 805 nm as indicated by different colors. Pigments with the corresponding in-vivo absorption maxima at the respective wavelength are shown in parentheses. c) Presence of genes associated with arsenic cycling and housekeeping genes (DNA gyrase (*gyrA*, *gyrB*)) in the metagenome of Pozo Bravo microbial mat. Shades of gray represent the number of gene copies (normalized as Transcripts Per Million, TPM^106^) from each microbial mat layer that were mapped to the metagenomic assembly, which was made using reads from all microbial mat layers. Almost as many gene copies are dedicated to arsenic cycling as for housekeeping tasks, suggesting intense metabolizing of arsenic species. d) Overview on the pathways of redox arsenic transformation detected in Pozo Bravo mats, based on previously reported functions of arsenic genes^20,108–110^. Genes were broadly classified in expressing transporters for arsenic uptake (blue) and export (red), enzymes involved in reduction of As(V) (orange), oxidation of As(III) (green), and production and export of organic arsenicals (pink). ArsC1 and ArsC2 are shown in gray since their presence could not be determined in the metagenome). Details and a list of abbreviations can be found in Table S1 and in the supplementary results Table 3.

To identify the key phototrophic groups across these layers, we combined hyperspectral imaging with metagenomic analysis. Despite recovering only three cyanobacterial bins – *Halothece* sp., ESFC-1 (*Spirulinaceae*) and RECH01 (*Elainellaceae*) – from the metagenomes (Extended Data Fig. 1, Suppl. Results 1), hyperspectral data confirmed that cyanobacteria were the dominant phototrophs in the mat. The green layer was dominated by Chl *a* (absorption maximum λmax≈690 nm) and phycocyanin (λmax≈625 nm) indicative of cyanobacteria^31–33^. This layer also included a zone dominated by BChl *a* (λmax≈810 nm), indicative of purple anoxygenic phototrophs^34^ (Fig. 1c). The red layer showed a clear absorption maximum at 715 nm, characteristic of BChl *e* in green anoxygenic bacteria like Chloroflexi^35^ or Chl *f* in cyanobacteria^36^.

Metagenome screening revealed a great variety of arsenic-related genes at levels comparable to housekeeping genes (Fig. 1c&d, Suppl. Results 2), suggesting arsenic metabolism is a core function of the mat community. Indeed, the total dissolved inorganic arsenic concentration in the water column was 6.9+/-0.4 µM, toxic to most life but substantially below levels in well-studied arsenic-rich ecosystems, such as Mono Lake^19,37^.

### Far-red light and dual-layer oxygenic photosynthesis

In-situ microsensor depth profiling revealed that OP occurred in two distinct mat layers. An upper oxygen concentration peak was observed at ∼1 mm depth (Fig. 2a&d, Extended data Fig. 2) in the green Chl *a*-dominated layer (Fig. 1b). An additional net OP maximum was observed at approximately 2-3 mm depth (Fig. 2a&d, Extended data Figs. 2, 3), coinciding with the red layer that exhibited an absorption peak characteristic of far-red pigments, consistent with either BChl *e* or Chl *f*. The detection of net oxygen production indicates a cyanobacterial origin, strongly suggesting Chl *f*, since anoxygenic phototrophs do not evolve oxygen. This spectral adaptation aligns with previously described strategies in cyanobacteria that extend photosynthesis into low-energy light environments^38–41^. These traits are believed to have evolved to support photosynthesis in shaded, low-energy environments – such as those within microbial mats – possibly already in the Precambrian^42^.

**Figure 2.**
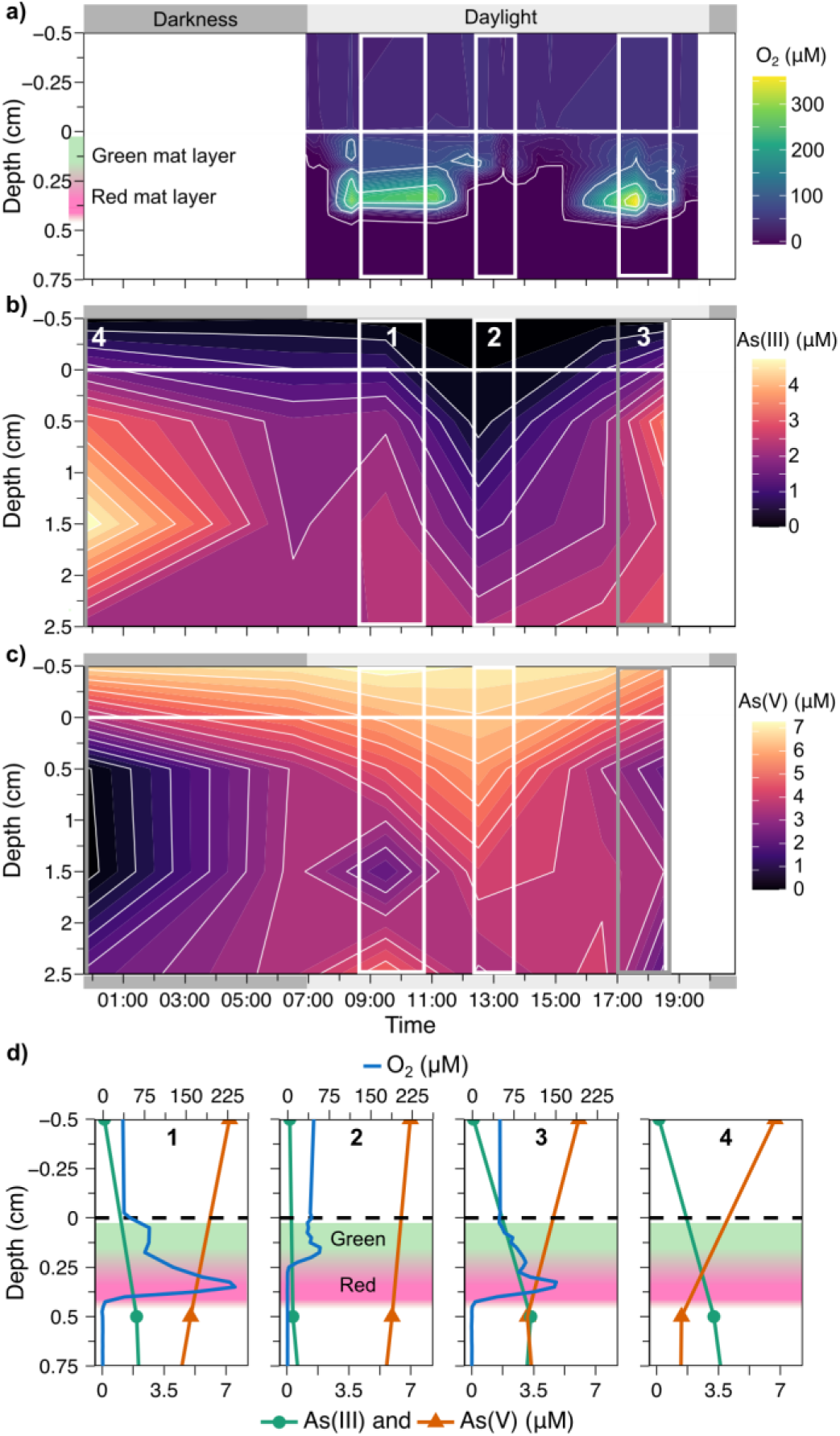
Diel dynamics of O_2_ production and arsenic speciation within the mats. Depth profiles of dissolved oxygen (a), and speciation of inorganic arsenic over a diel cycle in porewater from Pozo Bravo (b & c, n=1-3 per depth). The microbial mat surface is at 0 cm depth (horizontal white line in a)-c), black dotted line in d)). Profiles selected from various points in time are shown in d). The timepoints used for d) are indicated by rectangles in a)-c), following the numbering in b). Profile 4 represents a single point in time (00:30). Colored background represents the estimated placement of the green and red microbial mat layers (Fig. 1b). The chosen profiles exemplify the oxygen production during the morning (d1) and late afternoon (d3), with a period of interruption in between (d2). D4 shows a comparison of the dissolved arsenic speciation at night (00:30). The interruption in oxygen production coincides with a maximum in dissolved As(V). Net volumetric rates calculated based on the concentration depth profiles of oxygen measured in two replicate spots can be found in Extended data Fig. 3a, and depth integrated rates over a diel cycle in Extended data Fig. S3b.

### Diel arsenic cycling aligns with photosynthetic shutdown in deep mat layers

Microbial mats typically sustain continuous OP throughout daylight hours. However, in Pozo Bravo we observed an unusual diel pattern: OP declined in both phototrophic layers, and was even completely interrupted around midday in the deeper Chl *f*-dominated zone, despite ongoing light availability (Fig. 2a&d, Extended data Figs. 2, 3). This abrupt shutdown suggested that environmental stressors and/or metabolic transitions may regulate photosynthetic activity.

To investigate this, we monitored diel changes in key redox-sensitive compounds – arsenic, hydrogen sulfide, and iron – within the mat’s photic zone. Neither H₂S nor Fe(II) exhibited depth profiles or temporal dynamics consistent with phototrophic utilization (Supplementary Results 3). In contrast, arsenic speciation displayed pronounced, light-correlated shifts: As(III) dominated under dark conditions, while up to 98% of the dissolved arsenic pool was oxidized to arsenate (AsO ^3-^, As(V)) during the day (Fig. 2b,c). These dynamics were strongest within the top centimeter, where both cyanobacterial layers reside.

Strikingly, the peak in As(V) concentration coincided with the timing and depth of maximal OP inhibition in the deeper layer. This alignment suggests a potential causal link between arsenic redox transitions and photosynthetic regulation – specifically, that either arsenic itself or its light-driven oxidation may contribute to stress or otherwise disrupt OP, particularly in far-red light-adapted cyanobacteria.

### Reactive oxygen species link arsenic and light to photosynthetic inhibition

To study the effect of arsenic on OP, we performed ex-situ incubations of intact mat sections under varying light intensities, with and without added As(V). In arsenic-free conditions, oxygen production increased and stabilized under both low and high light, as expected for typical cyanobacterial mats (Extended Data Fig. 4). However, when 10 µM As(V) was added, oxygen production initially rose but then abruptly ceased after ∼30 minutes of high-light exposure (∼1000 µmol photons m^-2^ s^-2^). Oxygen production resumed when light intensity was reduced.

This response occurred independently of the time of day or duration of light exposure, indicating that OP suppression was not due to cumulative photoinhibition or diel rhythm, but instead resulted from a combined effect of high light and arsenic. Although most mat pieces did not exhibit the distinct double peaks observed in-situ, occasionally two net productivity maxima were observed. Notably, the response was most pronounced in deeper mat layers, mirroring in-situ patterns and suggesting that the Chl *f*-dominated community is particularly sensitive to this stressor combination.

To elucidate the mechanism behind OP inhibition, we simultaneously measured depth profiles of H_2_O_2_, a key ROS, under varying light and arsenic conditions (Fig. 3). Without arsenic, no net production of H_2_O_2_ was observed across light intensities. In As(V)-treated mats, H_2_O_2_ concentration within the photosynthetically active zone dramatically increased at 1000 µmol photons m^-2^ s^-1^. Upon the cessation of OP, H_2_O_2_ concentration dropped back to baseline levels. These dynamics suggest that ROS accumulation acts as a physiological signal or threshold, triggering the shutdown of OP in cyanobacteria to avoid further oxidative damage.

**Figure 3.**
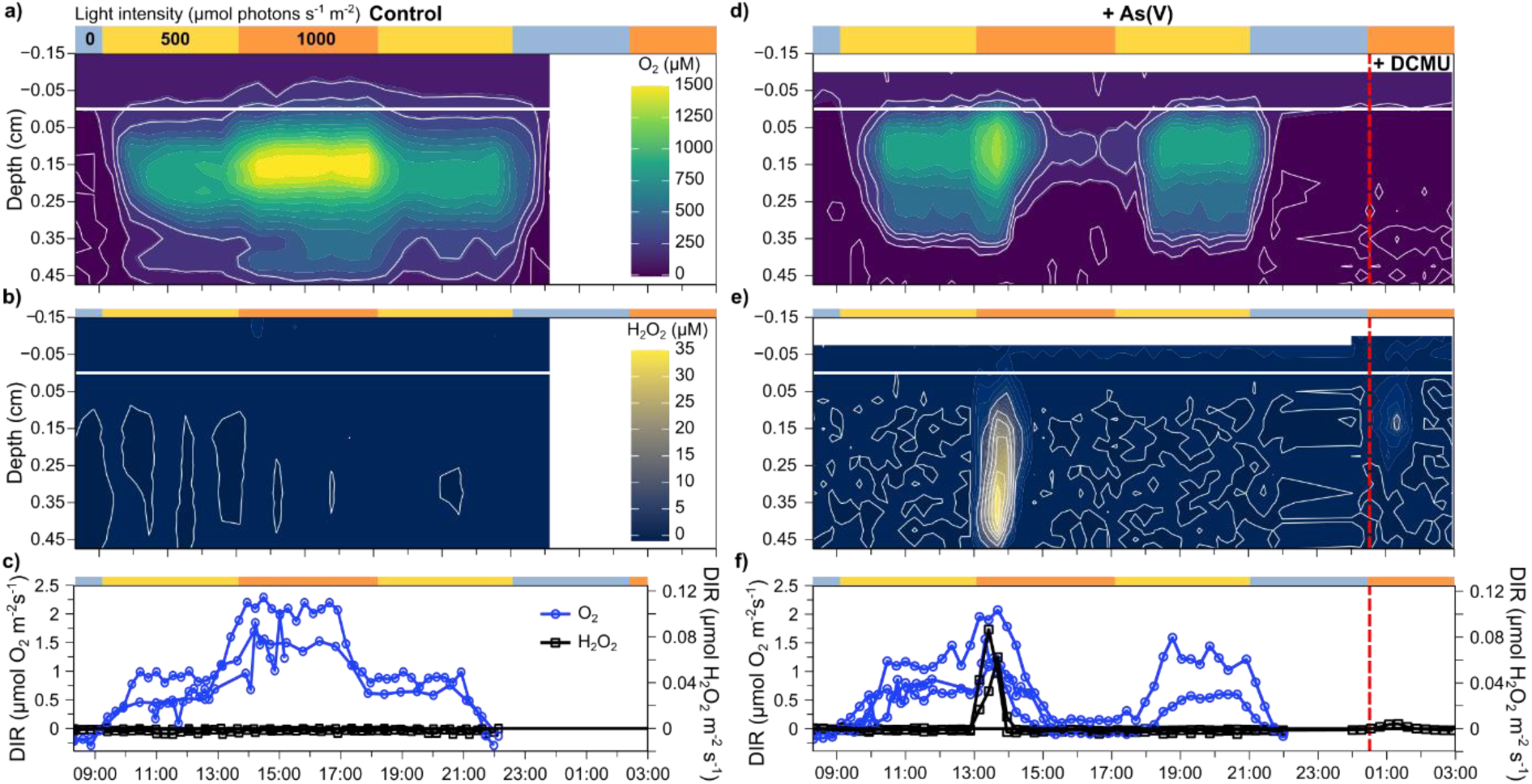
As(V) inhibits OP at high light and induces H_2_O_2_ evolution. O_2_ and H_2_O_2_ measurements of microbial mats before (a-c) and after As(V) addition (10 µM final concentration) (d-f)). Bars over each graph represent light conditions over time, as either darkness, low light (500 µmol photons m^-2^ s^-1^), or high light (1000 µmol photons m^-2^ s^-1^). Contour plots show concentration of O_2_ (a,d) or H_2_O_2_ (c,e) across the depth of the microbial mat over time. The solid white horizontal line represents the mat surface at depth 0. The vertical dashed line in d-f indicates time of DCMU addition, an inhibitor of oxygenic photosynthesis. c) and f) Depth-integrated rates of production (DIR) for O_2_ (n=3 pieces of mat) and H_2_O_2_ (n=2 pieces of mat).

Transcriptomic data from mats sampled in-situ across the diel cycle supported this interpretation (Fig. 5, Suppl. Results 4). Genes associated with oxidative stress and ROS defense – such as catalases, thioredoxins, and the phage shock protein *pspA* – were upregulated during mid-morning and around noon, coinciding with peak light and As(V) levels (Fig. 5b, Extended Data Fig. 7). Expression of key stress response genes like *pspA* and *hsp70* was detectable in the bin RECH01, consistent with membrane disturbance and ROS-mediated stress^43,44^.

While Pozo Bravo mats likely experience continuous oxidative pressure due to high irradiance and OP itself, the shift to net H_2_O_2_ accumulation suggests that sink capacity is overwhelmed under combined arsenic and light stress. Experimental addition of H_2_O_2_ confirmed that the mats can normally buffer external ROS: even at 200 µM H_2_O_2_, oxygen production remained unaffected and the compound was rapidly consumed in the upper layers (Extended Data Fig. 5), as also observed in other mat systems^14^. Thus, the observed H_2_O_2_ spike likely represents a breakdown of ROS homeostasis.

To identify possible sources of this ROS burst, we treated mats with both As(V) and DCMU, a PSII inhibitor that blocks OP. Even under these conditions, transient H_2_O_2_ production occurred at the oxic interface (Fig. 3f), indicating that OP is not the sole ROS source. Likely contributors include abiotic As(III) oxidation, light-driven ROS formation via iron photochemistry^45^, and singlet oxygen production during photosystem repair^46^. The reaction of As(III) with oxygen to form As(V) and H_2_O_2_ according to

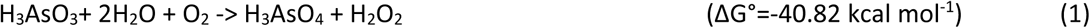

is thermodynamically favorable and may proceed both enzymatically and abiotically^47^.

Thus the combination of high light and dissolved arsenic induces local H_2_O_2_ accumulation that overwhelms the mat’s ROS-scavenging capacity, triggering a shutdown of OP – particularly in the deeper, far-red light-adapted cyanobacterial layer. This photophysiological response suggests a tightly regulated transition to an alternative metabolic mode under oxidative stress. Given the synchronous depletion of oxygen and accumulation of As(V), we hypothesized that the cyanobacteria switch to AP using As(III) as an alternative electron donor.

### Cyanobacteria perform arsenite-driven anoxygenic photosynthesis

To test whether cyanobacteria in Pozo Bravo mats can use As(III) for AP, we incubated mat pieces under various light wavelengths and inhibitor conditions over a diel cycle. The photosynthetic inhibitor DCMU was used to block PSII and thereby oxygenic activity, allowing us to detect only AP driven processes (Fig. 4, Extended data Fig. 6).

**Figure 4.**
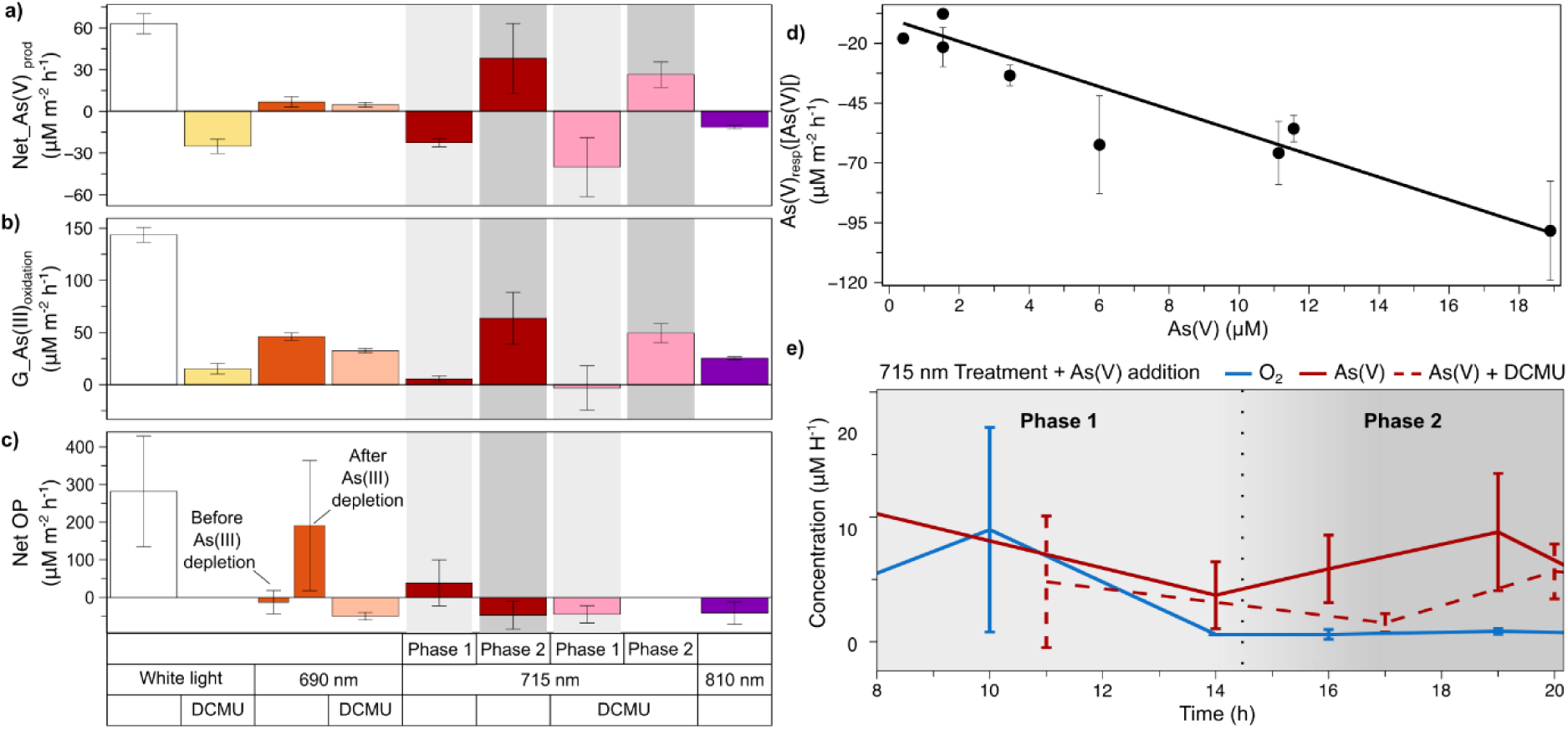
Rates of arsenic cycling dependent on light quality and photosynthesis. **a**) Net rates of As(V) production (Net_As(V)_prod_) in incubations of mat slurries with and without DCMU, under white light, light at λmax = 690 nm to target Chl *a*, λmax = 715 nm to target Chl *f*, and light at λ_max_ =810 nm to target BChl *a*. The gray bars overlaid in 715 nm represent phase 1 and phase 2 of As(V) production, as shown in (e). b) Rate of As(V) consumption, i.e. As(V) respiration, in the dark dependent on the concentration of As(V). c) Gross rates of As(III) oxidation (G_As(III)_oxidation_) calculated from (a) by adding the expected rate of As(V) removal (As(V)_resp_([As(V)]) from (b) according to Eq (1). d) Net rates of oxygenic photosynthesis in the same incubations as in (a) and (c). In e) an example for a time series of dissolved As(V) and O_2_ in batch incubations is shown, based on which the net rates of arsenic production in (a) and net OP in (d) were calculated. The slurry was initially exposed to ∼20 µM As(V) and, after a dark phase, exposed to light of λmax = 715 nm. Errors represent the standard deviation of concentration in separate vials (n=3). In phase 1, which represents the initial hours of exposure to light, net O_2_ production occurred and arsenite was not photosynthetically oxidized. In phase 2, after sustained exposure to light, O_2_ production ceased and As(III) was photosynthetically oxidized to As(V). A similar pattern was observed in the treatment with DCMU, which blocks OP. Namely, As(V) was reduced for an initial five hours (phase 1), after which As(III) was photosynthetically oxidized to As(V) (phase 2).

**Figure 5.**
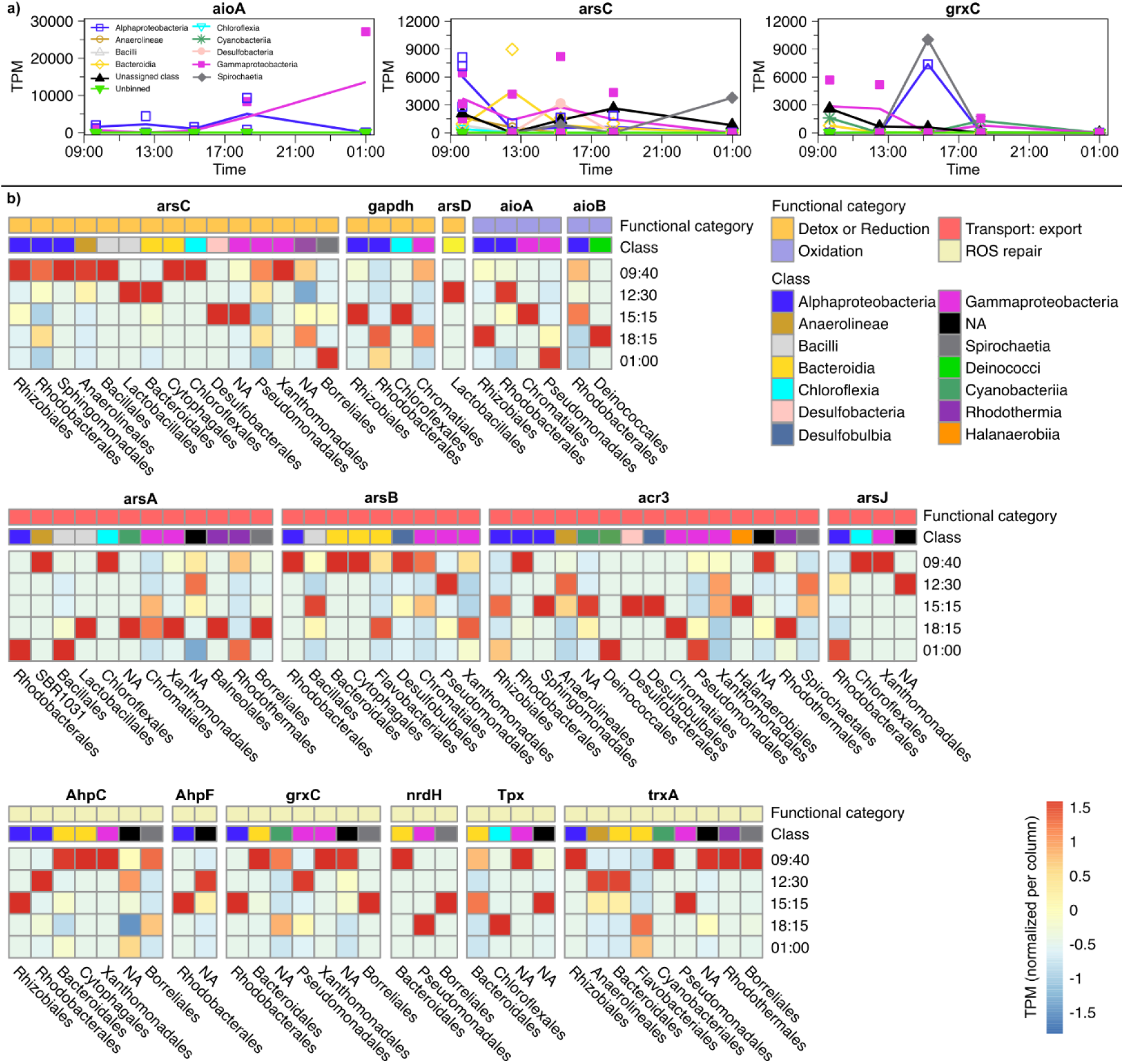
Transcriptional activity of arsenic-related genes across the Pozo Bravo microbial mat and selected taxa. a) Sum of *aioA*, *arsC*, and *grxC* transcription, in TPM, over the metatranscriptome of all layers of Pozo Bravo microbial mat. *aioA* is involved in arsenite oxidation, *arsC* is involved in arsenate reduction, and *grxC* may be part of arsenate reduction or reactive oxygen species response pathways. b) Transcription of genes related to arsenic cycling in selected taxa, shown as the sum of TPM per gene, normalized across all timepoints (i.e., per column). Normalization was performed due to the varying number of gene copies found per gene. Higher gene transcription is represented in red, lower transcription in blue. Transcription of *pufL* and *pufM*, which encode for photosynthetic reaction center II, can be found in the Supplementary Results (Fig. S2). An overview of transcription of genes in cyanobacterial bins can be found in Extended data Fig. 7.

In these incubations, we consistently observed net reduction of As(V) in the dark, likely by microbial respiration using As(V) as a terminal electron acceptor. Upon illumination, net reduction rates decreased, but in most cases, we did not observe net oxidation of As(III) – showing co-occurrence of light-driven As(III) oxidation and continued As(V) reduction. We thus estimated photosynthetic As(III) oxidation rates by correcting the net conversion for background As(V) respiration (Fig. 4a,d, See Methods).

These gross rate calculations revealed that light-driven As(III) oxidation occurred under all tested light conditions (Fig. 4). The highest rates were observed in white light without DCMU, consistent with aerobic oxidation using photosynthetically produced oxygen. However, gross oxidation persisted even in DCMU-treated samples – where OP was blocked – demonstrating true AP (Fig. 4b, Extended Data Fig. 6).

Light-quality experiments helped identify the phototrophs involved. At 810 nm, AP could be attributed to purple sulfur bacteria, as previously observed in cultures^19,37^ and in mats^22^. As(III) oxidation also occurred at 690 nm and 715 nm – wavelengths specific to cyanobacterial Chl *a* and Chl *f*, respectively. Crucially, gross oxidation was observed at these wavelengths in the presence of DCMU, strongly implicating cyanobacteria in As(III)-driven AP.

The transition dynamics between AP and OP differed markedly between exposures to 690 nm and 715 nm, suggesting that distinct cyanobacterial groups with different adaptations to As(III) were involved. The response at 690 nm followed a classic transition pattern, previously observed in cyanobacteria capable of sulfide-driven AP^10^: OP resumed only after As(III) was depleted. Thus Chl *a*-cyanobacteria initially performed AP and later switched to OP once the electron donor As(III) was exhausted (Extended Data Fig. 6d–i).

At 715 nm, a distinct biphasic pattern emerged, mirroring in-situ diel dynamics (Fig. 4a,c,e, Extended data Fig. 6l-q). During the first hours of illumination (phase 1), OP dominated and As(III) oxidation was low - an unexpected outcome given the high availability of electron donor for AP. Later (phase 2), OP ceased and As(V) production sharply increased, both in DCMU-treated and untreated samples (Fig. 4e). These patterns strongly support a delayed functional switch from OP to As(III)-driven AP by Chl *f*-containing cyanobacteria. This transition appears to require the combined presence of As(III), light, and oxygen, the latter accumulating only after prolonged illumination of the incubation vials.

Mechanistic insights into As(III)-driven AP was hindered by the low metatranscriptomic coverage of cyanobacterial bins. Although several arsenic-related stress and efflux genes (*arsA*, *arsB*, *acr3*^48^) were expressed (Extended Data Fig. 7), canonical As(III) oxidation genes (*aioAB*^49^) were not detected. This suggests that As(III) oxidation may involve divergent enzymes, or a fundamentally different biochemical pathway. Repeated attempts to isolate cyanobacterial strains capable of As(III)-driven photosynthesis were also unsuccessful, ultimately preventing mechanistic characterization of this process.

However, diel transcription of photosystem genes revealed dynamic regulation: PSI genes peaked in the morning, while PSII expression increased later in the day. The PSII core gene *psbA*, encoding D1, peaked shortly before OP resumed in the afternoon (Extended data Fig. 7b). Given that Chl *f* synthesis requires the replacement of D1 with a Chl *f*-specific isoform that inhibits PSII^50,51^, this expression pattern may suggest a regulatory mechanism by which cyanobacteria suppress OP and favor PSI-based AP under stress, then reassemble PSII to restore OP as conditions improve.

Together, these results provide strong indirect evidence that cyanobacteria in Pozo Bravo mats can perform As(III)-driven AP, particularly under far-red light and when OP is suppressed. This metabolic flexibility likely helps mitigate intracellular oxidative stress during periods of high light and arsenic exposure.

### A cryptic microbial arsenic cycle sustains phototrophic activity

The presence of As(III)-driven AP in the cyanobacteria-dominated mats of Laguna Pozo Bravo raises a key question: how is As(III) supplied during daylight hours, when in-situ measurements show that nearly all dissolved arsenic is present as As(V)? In this scenario, AP must rely on a continuous local supply of As(III). We therefore investigated how the broader microbial community sustains this process.

First, we examined metatranscriptomic profiles of arsenic redox cycling genes across the diel cycle. Focusing first on the oxidative side, transcripts of the canonical arsenite oxidase genes *aioA* and *aioB* were detected in several non-cyanobacterial taxa, including members of the Rhodobacterales (Alphaproteobacteria) and Chromatiales (Gammaproteobacteria), both including purple anoxygenic phototrophs and chemolithotrophs (Fig. 5b). These groups showed peak *aioAB* expression during daytime, coinciding with periods of active AP and elevated As(V) levels. This supports the notion that As(III) oxidation is a widespread phototrophic function within the mat.

To explore how As(III) remains available for As(III)-driven AP despite its low measured concentration, we examined the expression of genes involved in As(V) reduction, including *arsC*, *gapdh*, and *arsD*, which support both detoxification and respiratory pathways^52–54^ (Fig. 1). These genes, along with *arsA*, *arsB*, *acr3*, and *arsJ* (involved in As(III) efflux^48,52^), were transcribed throughout the diel cycle, with *arsC* expression peaking during the day (Fig. 5a). This suggests that As(III) is regenerated within the photic zone via continuous microbial As(V) reduction, effectively sustaining As(III)-dependent phototrophy even when As(V) dominates the bulk porewater.

Measured rates of As(V) reduction in incubation experiments supported this interpretation. Reduction rates scaled linearly with As(V) concentration up to ∼19 µM, showing no signs of saturation (Fig. 4d). These first-order kinetics are consistent with low-affinity reduction systems operating efficiently at ambient concentrations and suggest that As(III) production is dynamically coupled to its phototrophic oxidation. As a result, arsenic cycling within the mat is a cryptic redox loop, where As(V) is continually reduced and re-oxidized, enabling AP even under oxidizing conditions (Suppl. Results 5).

### A daily cycle of arsenic-driven photosynthesis and stress regulation

The microbial mats of Laguna Pozo Bravo operate as tightly regulated ecosystems in which phototrophic activity, arsenic redox cycling, and environmental stress responses are intricately linked. Over the course of a diel cycle, different microbial groups contribute to and are shaped by the dynamic chemistry of arsenic and light (Fig. 6).

**Figure 6.**
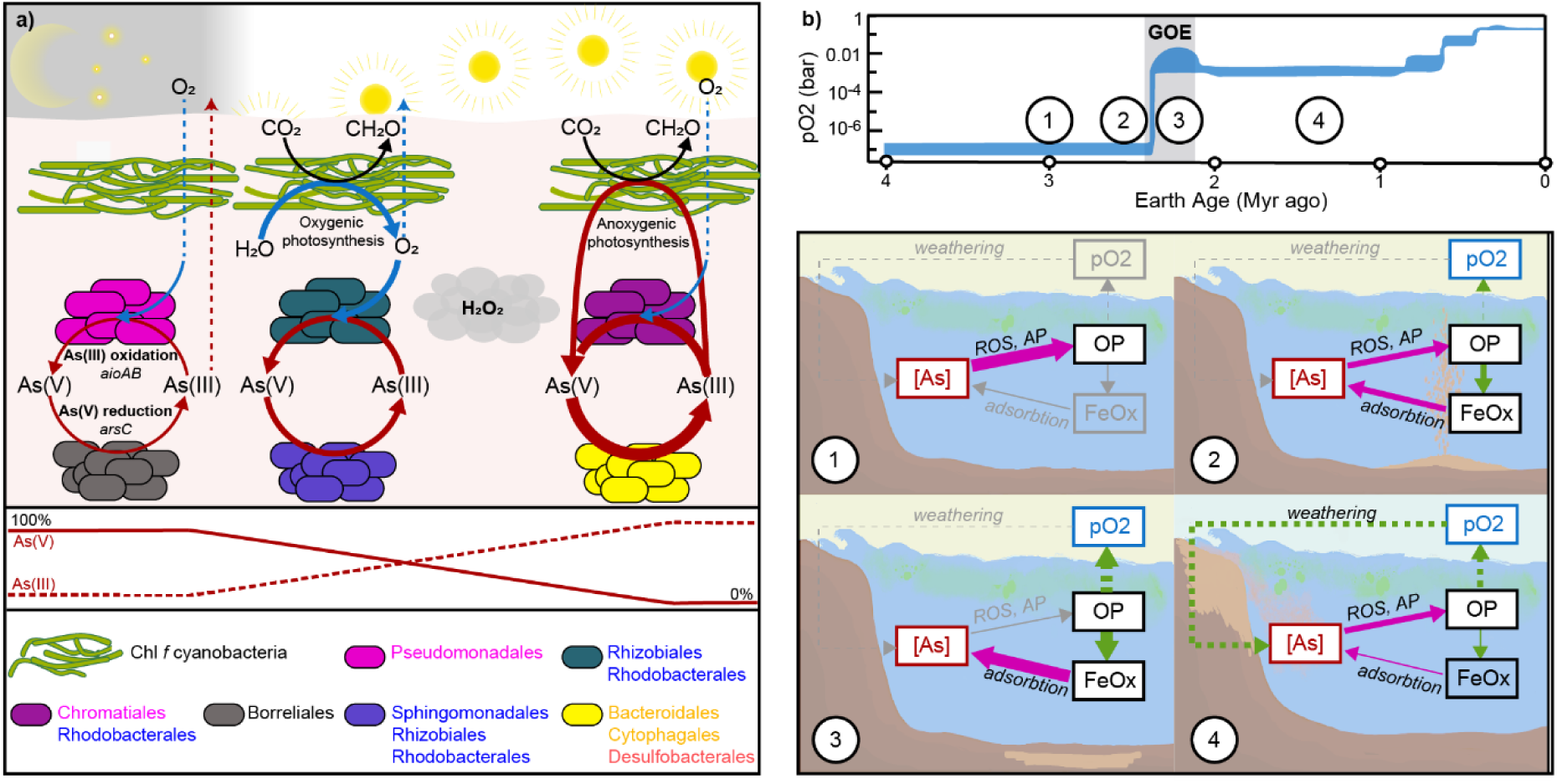
Proposed metabolic activities driving arsenic cycling through a diel cycle in the layers of Pozo Bravo microbial mats (a) and hypothesized feedback mechanisms between arsenic cycling and oxygenic photosynthesis in the Precambrian ocean (b). The mat model in (a) is based on pooling insights from hyperspectral imaging, process rate measurements and metatranscriptomic analyses. ROS production during the morning, afternoon and late afternoon is indicated with grey clouds. The conceptual model in (b) links these physiological responses to possible feedback mechanisms in the ancient ocean, with four numbered stages corresponding to the timeframes in the upper panel depicting Earth’s oxygenation history. 1. High As and ROS: OP is suppressed and cyanobacteria shift to AP, lowering net oxygen production. 2. OP recovery: Fe(II) oxidation by the photosynthetically produced oxygen yields Fe(III), which removes As(V) from the water column. 3. Low As and OP dominance: Reduced arsenic stress enables sustained OP and higher oxygen output. 4. Oxidative weathering: reintroduces As to oceans, reinstating OP suppression and tempering oxygen levels. Green arrows indicate enhancing effects, magenta arrows negative effects. Solid arrows: processes within the photic zone; dashed arrows: global-scale processes operating over geological timescales.

In the early morning, both cyanobacterial layers – those dominated by Chl *a* and Chl *f* – perform OP, producing oxygen and initiating the oxidation of As(III) to As(V). Anoxygenic phototrophs, including purple bacteria and possibly some Chl *a* cyanobacteria, also become active, performing AP using residual As(III) as an electron donor. In parallel, heterotrophic respiration with As(V) maintains a cryptic redox loop, releasing As(III) back into the mat and sustaining phototrophic arsenic cycling.

By midday, environmental stress peaks. High irradiance, and increasing aerobic As(III) oxidation to As(V) coincide with a measurable rise in H_2_O_2_. This oxidative stress triggers a shutdown of OP in the deeper, Chl *f*-dominated cyanobacteria, while surface-layer Chl *a* cyanobacteria maintain reduced OP rates, and possibly As(III)-driven AP sustained by the rising As(V) reduction rates. The deeper phototrophs transition completely to As(III)-driven AP.

In the late afternoon, light levels decrease and Chl *f* cyanobacteria gradually resume OP, completing the cycle. This synchronized sequence of OP, AP, and arsenic redox cycling supports a metabolically flexible community that responds dynamically to fluctuating light and chemical conditions. It highlights the ecological importance of arsenic – not just as a toxin or redox substrate, but as a regulating force in the coordination of microbial energy metabolism.

### Arsenic-mediated suppression of photosynthesis may have delayed planetary oxygenation

Although OP by cyanobacteria likely evolved during the Archean, it took hundreds of millions of years for atmospheric oxygen to accumulate substantially, culminating in the Great Oxidation Event (GOE) ∼2.5–2.3 billion years ago^55^. The reasons behind this lag remain debated, often in terms of ecological or geochemical limitations – such as nutrient scarcity, Fe(II)-driven AP competition, or inefficient organic carbon burial^7,8,56^. Our findings suggest an additional, previously unrecognized factor: arsenic could have directly suppressed OP in early microbial ecosystems.

In modern Precambrian analogs at Pozo Bravo, we observed that cyanobacteria dynamically transitioned from OP to As(III)-driven AP. This shift was not driven by substrate depletion or light limitation, and appears to reflect a physiological response to oxidative stress rather than a passive metabolic fallback. When ROS levels rise cyanobacteria suppress OP – but instead of halting phototrophy, they re-route electron flow to perform AP with As(III) as the electron donor. This strikingly adaptive strategy avoids intracellular ROS from PSII activity and extracellular ROS generated by As(III) oxidation in the presence of oxygen (Eq. 1), while maintaining energy conservation. Such plasticity likely conferred a major advantage in early Earth environments, where UV stress, metal toxicity, and redox fluctuations were pervasive, positioning As(III)-driven AP as a purposeful stress-avoidance strategy rather than an evolutionary relic.

This strategy comes at a cost for the ecosystem’s net oxygen budget – as AP does not release oxygen. Notably, the mitigation of OP does not require high arsenic levels, because tight redox recycling maintains As(III) availability. Such cryptic cycling may have enabled arsenic to act as a persistent, structurally embedded electron donor in early microbial ecosystems. Thus, even in low-arsenic environments cyanobacteria could have repeatedly faced conditions where OP was inhibited and replaced by AP, limiting cumulative oxygen output.

Our findings show that arsenic did not merely act as a passive redox substrate in early environments, but may have actively modulated the pace of oxygenation through physiological inhibition of OP. Namely, OP may have initially inhibited itself: the oxygen it produced aggravated oxidative stress in arsenic-rich environments. However, this negative feedback may have transitioned into a positive loop, in which rising levels of oxygen ultimately promoted the widespread oxidation of both As(III) and Fe(II)^57,58^. In response to rising As(V), genomic evidence indicates a vast expansion of arsenic metabolism genes across bacteria^59^. Yet, As(V) is known to strongly sorb onto freshly precipitated Fe(III) oxides, effectively removing arsenic from the water column^17,60^. Here we show that this would have decreased arsenic-induced oxidative stress and relieved inhibition of OP, allowing photosynthetic activity – and oxygen accumulation – to increase further. Such a feedback loop may have contributed to the eventual crossing of the critical tipping point for oxygenation during the GOE.

Following the GOE, Fe(II) was largely depleted from the oceans, reducing Fe(III) oxide formation and As(V) scavenging^61^. During the mid-Proterozoic “boring billion,” continental oxidative weathering, persistent anoxia and the expansion of sulfidic conditions further hindered arsenic sequestration^17,57^. Consequently, arsenic could have re-emerged as a physiological stressor, periodically constraining OP and helping maintain low oxygen levels during much of the Proterozoic.

Crucially, arsenic may have had a suppressive effect on OP itself, while simultaneously stimulating AP. This shift would have maintained high primary productivity and organic carbon burial, but without the corresponding oxygen release. By decoupling carbon burial from oxygen accumulation, this mechanism may help explain apparent discrepancies between signals for productivity and oxygen levels in the rock record after the GOE^57,62,63^. Arsenic cycling thus emerges as a potentially critical factor in modulating biogeochemical feedbacks during Earth’s oxygenation, yet remains underexplored in most current models.

## Methods

### Field measurements and sampling

Field measurements and sampling were conducted in Laguna Pozo Bravo (25° 30’ 58” S, 67° 34’ 42” W), in the Salar de Antofalla, Argentina in late February 2019 and January 2023 at an elevation of 3330m. To assess diel dynamics of oxygen and H_2_S within the mat, we continuously measured depth profiles of concentration on two consecutive days, in two spots of the mat. The microsensors (H_2_S and Clark-type O_2_ with <50 µM tip width) were constructed, calibrated and used as previously described^64–67^. Vertical profiles were measured simultaneously by attaching the microsensors to a multi-sensor holder, yielding a distance between tips of ∼0.5 cm. Electricity was provided by a portable generator. Using O_2_ microsensor measurements, we calculated the net rates of OP as the difference in the fluxes into and out of the two photosynthetically active zones.

In parallel with the microsensor measurements, subsamples of the mat were regularly taken for porewater extraction. Cores for porewater extraction (2.5 cm inner diameter) were sampled in triplicate every three hours from 1 am until 6 pm. Porewater was immediately extracted from the cores (2.5 cm ID, up to 6 cm depth) at 1 cm intervals using rhizones (Rhizosphere Research Products, NL). An aliquot of 500 µL porewater was instantly frozen with liquid nitrogen and stored and transported in a dry shipper until arsenic speciation analysis as previously described^68^. Briefly, arsenic speciation was determined by high performance liquid chromatography (HPLC; 1260 Infinity II bio inert, Agilent) using a PRP-X100 column (Hamilton 5 µm, 10 mM NH_4_NO_3_, 10 mM NH_4_H_2_PO_4_, and 500 mg/L Na_2_-EDTA at a flow rate of 1.0 mL/min and 25 μL injection volume) coupled to ICP-MS/MS (8900 Triple Quadrupole, Agilent). Arsenic was detected in MS/MS mode using oxygen as reaction cell gas (AsO^+^, m/z 75 -> 91). Retention times of arsenite and arsenate were determined using individual standards. Samples stabilized in 2% HNO_3_ were analyzed for total As concentrations by ICP-MS/MS using Rhodium (Rh) as an internal standard (AsO^+^, m/z 75 -> 91; Rh^+^ 103 -> 103).

Additional sediment cores for meta-omics (2.5 cm inner diameter, up to 4 cm depth) were sampled in triplicates every three hours. The microbial mat was rapidly and carefully cut into four pieces per core that were instantly frozen in liquid nitrogen for metagenomic and metatranscriptomic analyses.

A few additional cores were frozen intact for hyperspectral imaging. The core was sliced in half and embedded with resin (EPO-TEK® MED-301-2FL) following the manufacturer’s instructions. The cross section of the core was then scanned using a Resonon Pika II hyperspectral camera, as described in Chennu et al. (2013)^69^. Scans were used to record radiance images of 0.2 mm per pixel in 4 bands of 1 nm at 675, 715 and 805 nm. Reflectance images were derived by normalizing the radiance spectra of all pixels to the average radiance of a standard reference board in each image.

Further, 10 x 5 cm pieces of mat with ∼0.5 cm underlying sediment were stored in plastic containers for transport and maintained in mesocosms at ∼700 µmol photons m^-2^ s^-1^ underneath a water column with a salinity of ∼137 g L^-1^ (TropicMarine artificial saltwater), constantly circulated to maintain a temperature of 21°C, until ex-situ measurements.

Temperature of air and water above the mat were recorded using HOBO loggers (Onset Computer Corp., MA). Measurements of light intensity failed due to rapid oversaturation of the sensor signal in the morning. Occasionally, pH measurements in the water column were taken using a Mettler Toledo pH meter. Salinity was measured with an Atago refractometer.

### Assessment of diel H_2_O_2_ dynamics

To assess the effect of light and arsenic on net O_2_ production and H_2_O_2_ dynamics in the mat, we subsampled microbial mat pieces (∼ 2.5×2.5 cm) from the laboratory mesocosms and continuously measured O_2_ and H_2_O_2_ depth profiles during simulated diel light cycles, in the absence and presence of dissolved arsenic. Mat pieces were cut and fixed with low-melting agarose (1%) to the bottom of glass dishes. Circular flow was established with the same hypersaline water (TropicMarine artificial saltwater approximately 137 g L^-1^) used for regular maintenance of the mats, and adjusted using a peristaltic pump (Minipuls 3 Abimed, Gilson). Diel cycles were simulated using halogen light sources (KL 2500 and KL 1500, Schott). Microsensors (H_2_O_2_, and O_2_ with 80-120 µM tip diameter) were constructed, calibrated and used as previously described^67,70,71^. Concentration-depth profiles were performed under three light conditions: dark, 500 and 1000 µmol photons m^-2^ s^-1^. First, profiles were measured without supplementing As(V), assuming that most arsenic had been washed out after four months of lab maintenance. Then As(V) was added to the overlying water column to a final concentration approximately 10 µM, mimicking the environmental As(V) maximum. After As(V) addition, mats were left undisturbed overnight to enable the diffusion of arsenic into the mat, and diel profiling began in the morning. Diel profiling was also performed in As(V)-supplemented mats after addition of DCMU, an inhibitor of OP, and after autoclaving the mat as an abiotic control.

Depth-integrated rates of net O_2_ and H_2_O_2_ production were calculated from the concentration depth profiles by considering Fick’s second law of diffusion. First, local volumetric net rates were calculated as the second derivative of concentration multiplied by the diffusion coefficient corrected for temperature and salinity. Volumetric depth rates were then integrated over the uppermost 5 mm of the mat.

### Assessment of arsenic cycling rates in batch incubations

To assess the dependency of light quality on arsenic cycling, small subsamples of microbial mats were incubated under As(V), As(III), and LED lights targeting distinct groups of phototrophs. Hypersaline water (TropicMarine artificial saltwater ∼137 g L^-1^) was deoxygenated by bubbling with helium, vigorously shaking, then removing gas bubbles, repeated five times. Subsamples cut from microbial mats (∼200 mg) using 8 mm width cores (surface area 0.5 cm^2^) were inserted into 3 mL gastight glass vials with a septum cap (exetainers, Labco) which were later filled completely with deoxygenated hypersaline saltwater. Then they were placed into containers on shakers, with constant flow of temperature-regulated water. The incubations were subjected to 8-hour cycles of light with various different spectral quality, followed by 8 hours of darkness. Controls were placed under constant darkness. All light treatments were repeated with the addition of 5 µL DCMU solution (5 mM) to inhibit oxygenic photosynthesis.

The light treatments were chosen at specific wavelengths to target different groups of phototrophs. LEDs with an emission maximum (λ_max_) at 690 nm (hexagonal high-power LEDs from Roithner Lasertechnik, Vienna; Supp. Fig. S18) were used to specifically target Chl *a*, i.e., the light harvesting pigment of most cyanobacteria for photosystem I and the central pigment in the reaction center of both photosystems. Since an unexpected absorption maximum was found at 715 nm in the lower cyanobacterial layer, we also used LEDs in the far-red range of the spectrum with λ_max_ = 715 nm to target cyanobacterial Chl *f*, which can likely act as antenna in both photosystems^41,72^. Additionally, we used λ_max_ = 810 to target BChl *a* of purple anoxygenic bacteria, and full-spectrum white light to target all phototrophs.

Every three hours, three incubation vials per treatment were removed for measurements. First, oxygen in each exetainer was measured with a microsensor, then 2 mL of the overlying water were filtered. From the filtered water, 1 mL was fixed with 4 mL 2% zinc acetate for total dissolved sulfide measurements, and 1 mL was fixed with 15.53 µL fixing solution and flash frozen for arsenic measurements. Fixing solution for arsenic measurements was made by mixing 27 mL of concentrated HNO_3_ diluted 1:2 with ultrapure water and 100 mL fresh DTPA solution. The DTPA solution was made by mixing 6.75 mL ultrapure water, 6.75 mL concentrated HNO_3_ and 50 mL of 40% premade DTPA*Na_5_ solution (Sigma Aldrich, CAS 140-01-2). Sulfide was later measured as previously described^10,73^.

Arsenic was measured colorimetrically based on Castillejos Sepúlveda et al. (2022)^68^, with modifications to calculations due to the high salinity and absence of phosphate. Specifically, instead of 1.15 mM KIO_3_, KMnO_4_ at a final concentration of 6.9 mM was used as the oxidant to convert As(III) to As(V). This is because oxidation by KIO_3_ was shown to be very inefficient at high salinity. As the analysis protocol from Castillejos Sepúlveda et al. (2022)^68^ could also not be completely adapted for hypersaline samples, we optimized the spectral analysis for the specific samples used here. The approach was rigorously simplified because dissolved inorganic phosphate concentration, an interfering solute, was below detection limit throughout. Absorption in the low-acidity reagent, which is proportional to both

As(V) and phosphate (not present) was measured at 648 nm and 760 nm. Concentration of As(V) in the samples was determined based on the difference between absorption at the wavelength, i.e. the spectral index, and a calibration series in the same hypersaline water. To assess the concentration of As(III) absorption at the same wavelengths was determined after oxidant addition and a reaction time of 3.5 h. The difference between the spectral index before and after addition was used to calculate As(III) based on calibrations with As(III) standards in hypersaline water with the same composition as the samples.

Incubations were initiated by the addition of As(V) or As(III) to enable differentiation between oxidation and reduction rates. As total dissolved arsenic remained relatively constant in all incubations, we calculated rates of both As(III) oxidation and As(V) reduction based only on As(V) dynamics, which was favorable due to the higher accuracy of the colorimetric method for As(V)^68^.

Given that As(V) respiration and photosynthetic As(III) oxidation likely co-occur, we conducted a detailed analysis of As(V) consumption (i.e., respiration) rates to enable the calculation of photosynthetic gross rates from net rates. Rates of As(V) consumption in the dark were linearly dependent on As(V) concentration across the complete measured range (Fig. 4b). Assuming that As(V) respiration rates were unaffected by light and solely determined by the As(V) concentration, we were able to calculate light-driven gross As(III) oxidation as

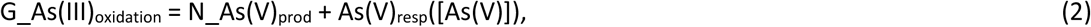

where G_As(III)_ox_ is the gross rate of As(III) oxidation (i.e. gross As(V) production, Fig. 4b), N_As(V)_prod_ is the net rate of As(V) production, (i.e. net As(III) oxidation), and As(V)_resp_([As(V)]) is the rate of light-independent As(V) consumption as a function of concentration derived from the linear regression in Fig. 4d.

### Metagenomics

A piece of microbial mat collected in situ at 12:30 pm, following the procedure described above, was sliced into the four distinct layers (crust, green, red, brown) and sent to the Max Planck Genome Center Cologne, Germany (https://mpgc.mpipz.mpg.de/home/) for processing. DNA extraction was performed using the DNeasy PowerSoil Pro Kit (Qiagen). An Illumina-compatible library was prepared with NEB Ultra II DNA kit (NEB) and each library was shotgun sequenced with Illumina HiSeq3000 (2 x 150 bp, paired-end read mode). The crust and brown layers were sequenced at a depth of 6.8 million reads, while the green and red layers were sequenced at a depth of 34 million reads to target cyanobacteria, which are notoriously difficult to sequence^74^.

Raw sequences were quality-trimmed with Trimmomatic v.0.38^75^, and assembled with MEGAHIT v.1.2.9^76^. Initially, a co-assembly of all samples was used, but due to low cyanobacterial read depth, a second assembly was prepared using only reads from the green layer of the microbial mat. Both assemblies were later processed using the same workflow, but the second assembly was only used for analysis of cyanobacteria.

Samples were mapped onto the assembly with Bowtie2 v.2.4.5^77^. Gene calling was performed with Prodigal v.2.6.3^78^, and the resulting gene calls were annotated in anvio v.7.1^79^ using the KEGG KOfam v.95.0^80^ and COG v.2020^81^ databases with DIAMOND v.0.9.14^82^ search. Interproscan v.78^83,84^ was used for further annotations with the following databases: CATH-Gene3D v.4.2.0^85^, CDD v.3.17^86^, HAMAP v.2020_01^87^, PANTHER v.14.1^88^, Pfam v.32.0^89^, PIRSF v.3.02^90^, PRINTS v.42.0^91^, PROSITE Patterns and PROSITE Profiles v.2019_11^92^, SMART v.7.1^93^, SFLD v.4^94^, SUPERFAMILY v.1.75^95^ and TIGRFAMs v.15.0^96^. Genes related to arsenic, photosynthesis, glutaredoxins, and thioredoxins were manually selected and annotated based on consensus from the automatically-generated annotations, using BLASTx v.2.2.31^97^ to resolve discrepancies.

Metagenomic analyses were initially focused on genes that could result in redox changes, before binning was performed to assign genes to distinct bacterial taxa. Automatic binning was done with MetaBAT2 v.2.15^98^ on the large co-assembly and MaxBin2 2.2.7^99^ on the second assembly. In both cases, bins were manually refined in anvio. Taxonomy was assigned to bins using GTDB-TK v.95.0^100,101^ within anvio. The relative abundance of each bin was calculated using coverM v. 0.4.0^102^ and plotted using ggplot2^103^ in R v. 4.0.3^104^.

### Metatranscriptomics

Microbial mat samples collected following the procedure described above in situ at 1:00 am, 9:40 am, 12:30 pm, 3:15 pm, and 6:30 pm were processed at the Max Planck Genome Center Cologne, Germany. RNA was isolated using Quick-DNA/RNA Miniprep Plus Kit (Zymo Research). Next, an Illumina-compatible library was prepared with the kit Universal Prokaryotic RNA-Seq incl. Prokaryotic AnyDeplete for rRNA depletion (Tecan Genomics). Each sample was shotgun-sequenced with Illumina HiSeq3000 (2×150 bp, paired-end read mode) at depths ranging from 35.5 million to 51 million reads. Raw sequences were quality trimmed with Trimmomatic, then mapped onto the metagenomic assemblies with BBmap v38.73^105^, and used featureCounts from Rsubread v. 1.22.2 to obtain counts of each gene per metatranscriptomic sample. To compare gene transcription across samples, counts of each gene call were normalized by calculating Transcripts Per Million (TPM) as previously described^106^. The TPM of all gene calls annotated with the same function (i.e.-the same gene name) were summed, and used to make heatmaps with Pheatmap v.1.0.12^107^ in R.

## Data availability

Data has been submitted to the EMBL Nucleotide Sequence Database (ENA). Metagenomic and metatranscriptomic raw reads, as well as cyanobacterial metagenome-assembled genomes can be found under project number PRJEB41764. Metagenome-assembled genomes from the co-assembly can be found under project number PRJEB73824.

## Supporting information

Supplementary Results

## Acknowledgements

We would like to thank Secretaría de Medio Ambiente de la Provincia de Catamarca for providing the research permit (Expte.EX-2022-02222431-CAT-DPB#SEAS), and Secretaría de Política Ambiental en Recursos Naturales, Ministerio de Ambiente y Desarrollo Sostenible de la Nación Argentina for providing the certificate of compliance (Title: IF-2023-65966642-APN-SPARN#MAD, UId: ABSCH-IRCC-AR-264943-1) and the export certificate for genetic resources (CE-2023-69835755-APN-SPARN#MAD). We would like to express our gratitude to Luis Ahumada for his assistance during the field campaigns, and the native communities of Antofalla, El Peñón, and Antofagasta de la Sierra for their valuable support. We the Max Planck-Genome-centre Cologne (http://mpgc.mpipz.mpg.de/home/) for performing DNA and RNA extractions and sequencing, in this study. We are thankful to Gert Bange, Pieter Visscher, Christophe Dupraz, Pascal Philippot and Brendan Burns for fruitful discussions. The authors also thank the laboratory technicians in the Microsensor working group, the mechanical workshop in MPI Bremen, David Benito Merino, Marit van Erk, Britta Planer-Friedrich, Martina Alisch, and Boran Kartal for their help with labwork, advice, and laboratory equipment.

## Author contributions

J.M.K. and A.C.S. conceptualized the study. J.M.K. supervised the project. A.C.S. and J.M.K. led the experiments and wrote the first draft. Incubation experiments were designed and performed by A.C.S., J.M.K., W.M., and E.M. Ex-situ microsensor measurements were conducted by K.S. and J.M.K. Arsenic speciation analyses were carried out by C.K., J.M.K., and A.C.S. Meta-omics analyses were performed by D.R.S., F.A.V., and A.C.S. A.C. conducted hyperspectral imaging. Fieldwork was carried out by A.C.S., J.M.K., D.D-W., F.A.V., E.M., and M.E.F. Figures were prepared by A.C.S., J.M.K., and D.D-W. Resources and materials were provided by M.E.F., D.d.B., and J.M.K. All authors contributed to and approved the final manuscript.

---

**Extended data Figure 1.**
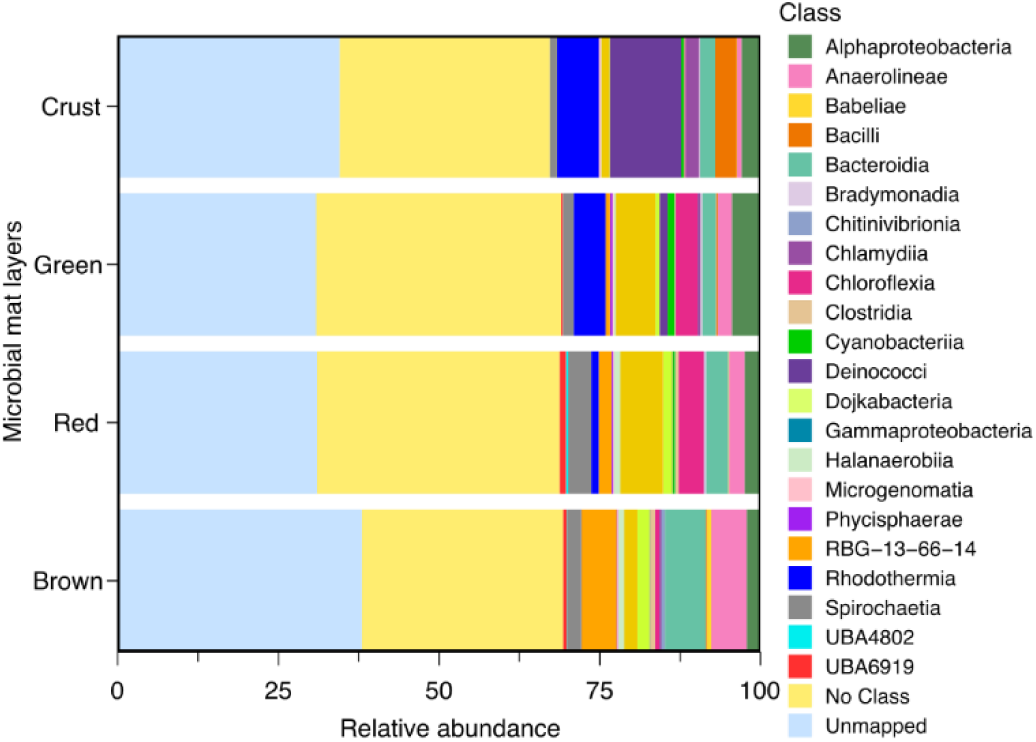
Taxonomic profile of phyla from metagenomic bins in a Pozo Bravo microbial mat. The bins were selected from a metagenomic assembly made using reads from all vertically-stratified layers (crust, green, red, brown). Only bins with over 55% completeness and less than 10% redundancy are shown (Supplementary Table S1). Relative abundance was obtained by mapping the reads from each individual layer metagenome to the total set of bins. Results were summarised by grouping bins by phylum.

**Extended data Figure 2.**
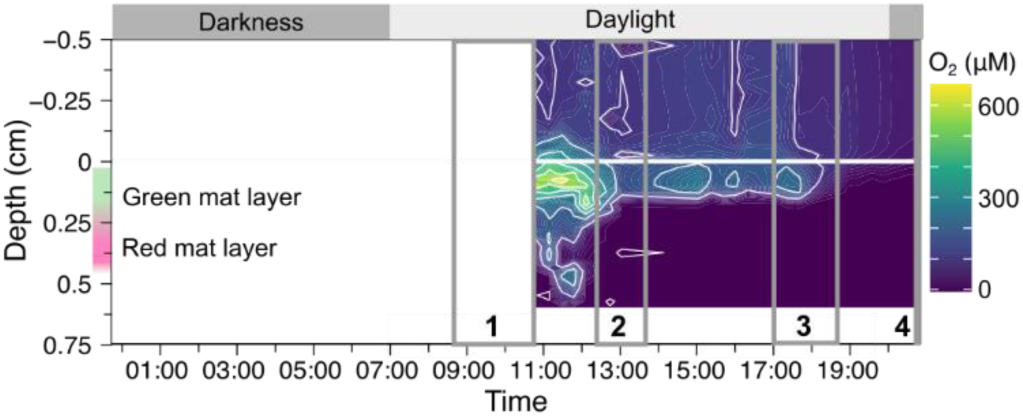
Depth profiles of dissolved oxygen in Pozo Bravo microbial mat, replicate of Fig. 2a and corresponding to “mat b” in Extended data Fig. 3. The microsensor profiles shown in Extended data Fig. 3 were selected from times (1-4), as shown by the gray boxes with black numbers.

**Extended data Figure 3.**
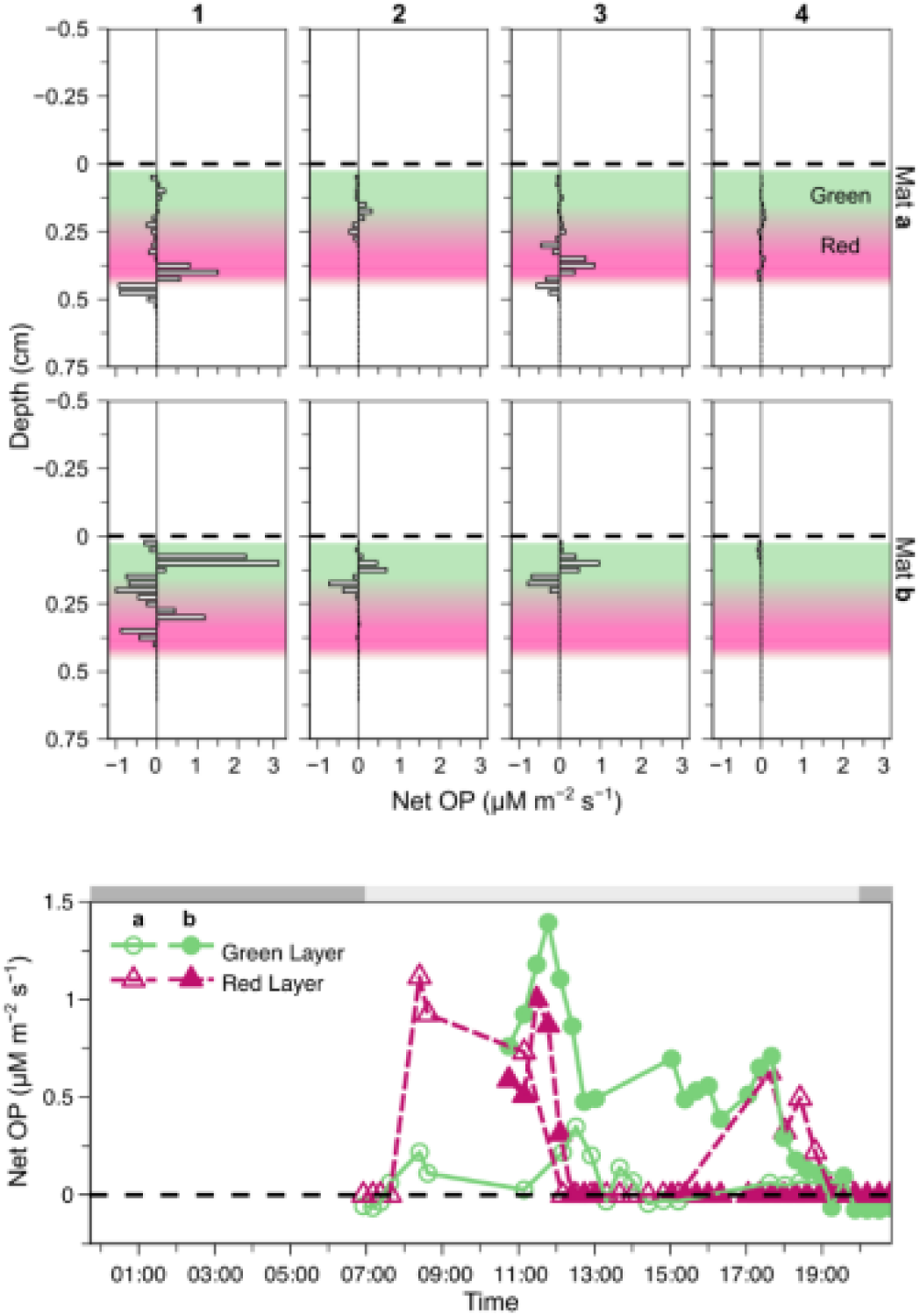
Net volumetric rates of oxygenic photosynthesis. In a), examples for local volumetric rates are shown through depth, calculated from the oxygen profiles shown in Extended data Fig 2 and in Fig. 2a in the main text, at the times specified by 1-4 in Fig 2. Colored bars indicate the estimated location of the green and red microbial mat layers. In b), the corresponding depth-integrated rates of net OP over a diel cycle in the green and red layers are shown.

**Extended data Fig. 4.**
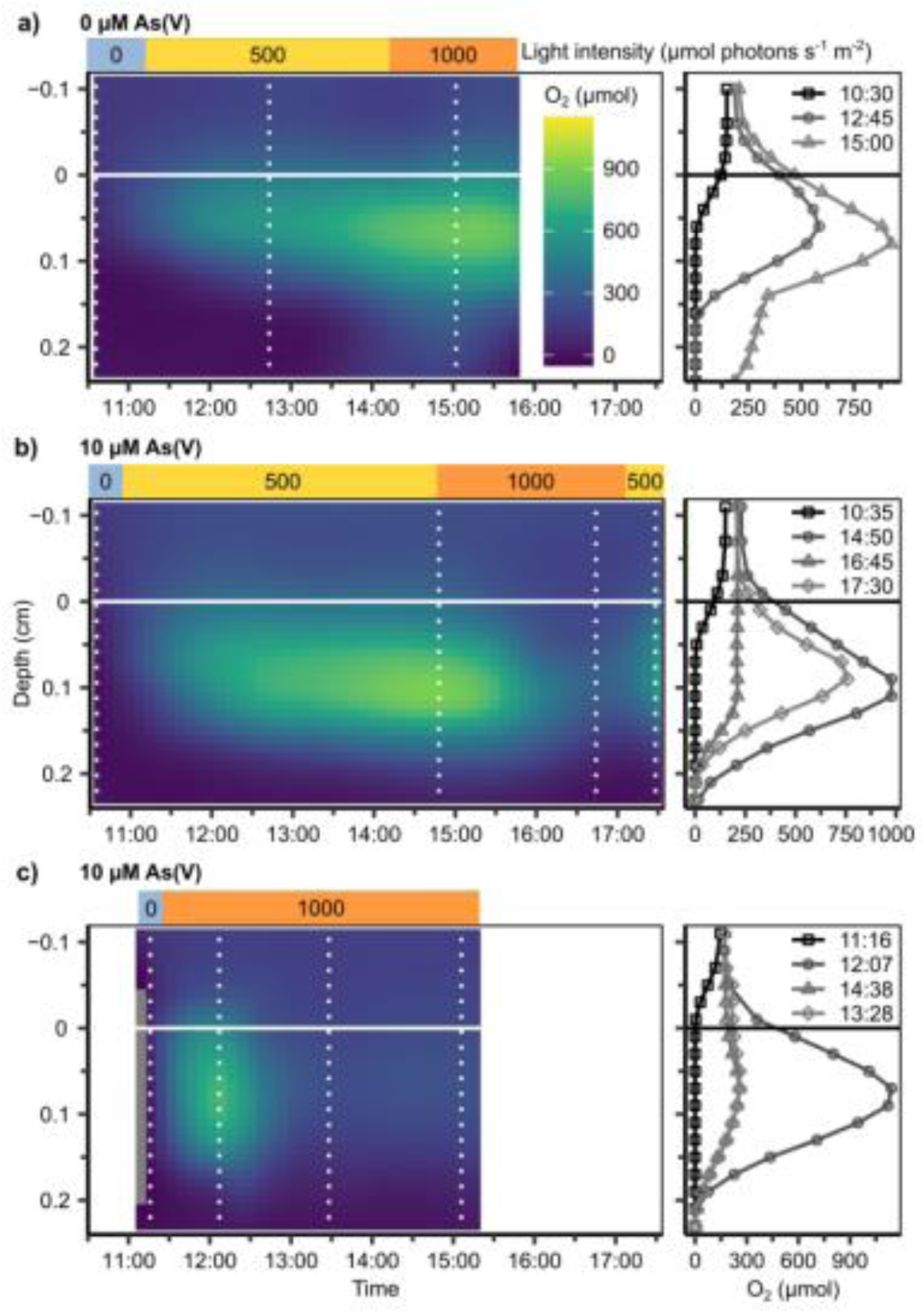
Microsensor measurements of Pozo Bravo microbial mat incubations under two light conditions (500 and 1000 µmol photons m-2 s-1), with and without 10 µM As(V) addition. Selected profiles, indicated by vertical dotted lines, are shown on the right panels. Microbial mat surface is indicated by horizontal lines at depth 0. a) and b) were obtained from the same piece of microbial mat, while c) was measured in a separate mat sample.

**Extended data Figure 5.**
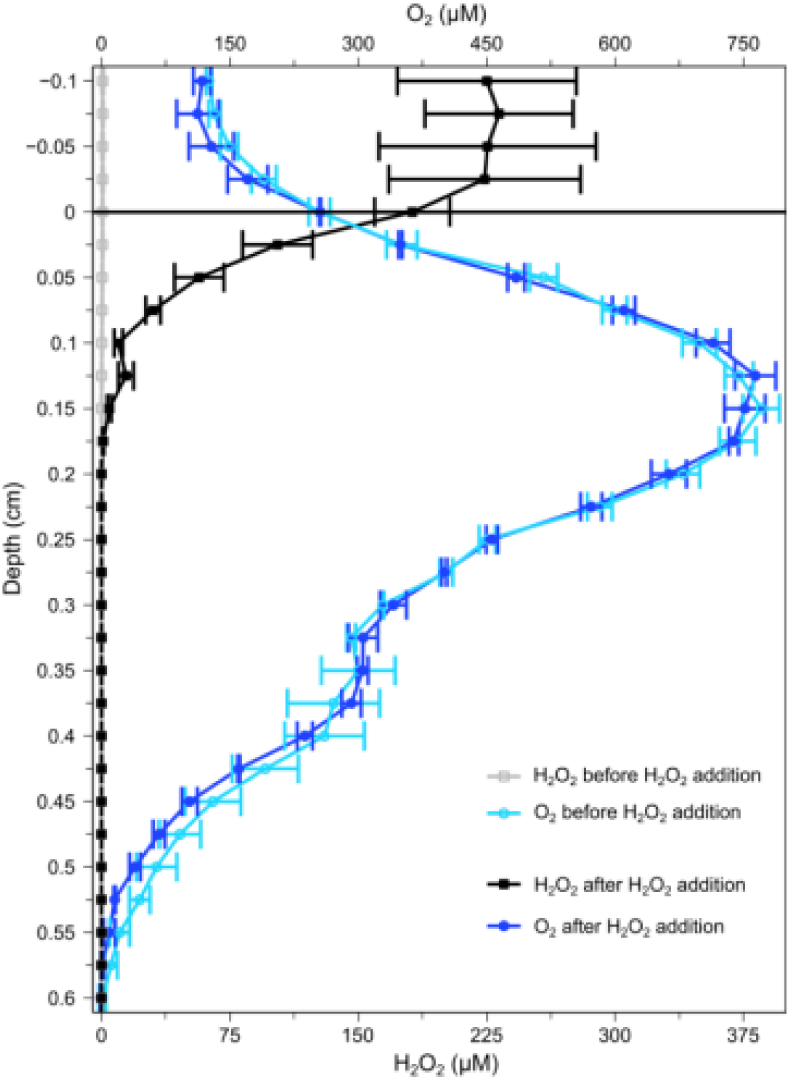
Ex situ H_2_O_2_ and O_2_ microsensor depth profiles of Pozo Bravo microbial mat, taken before and after the addition of H_2_O_2_ to the water column (n=3).

**Extended data Fig. 6.**
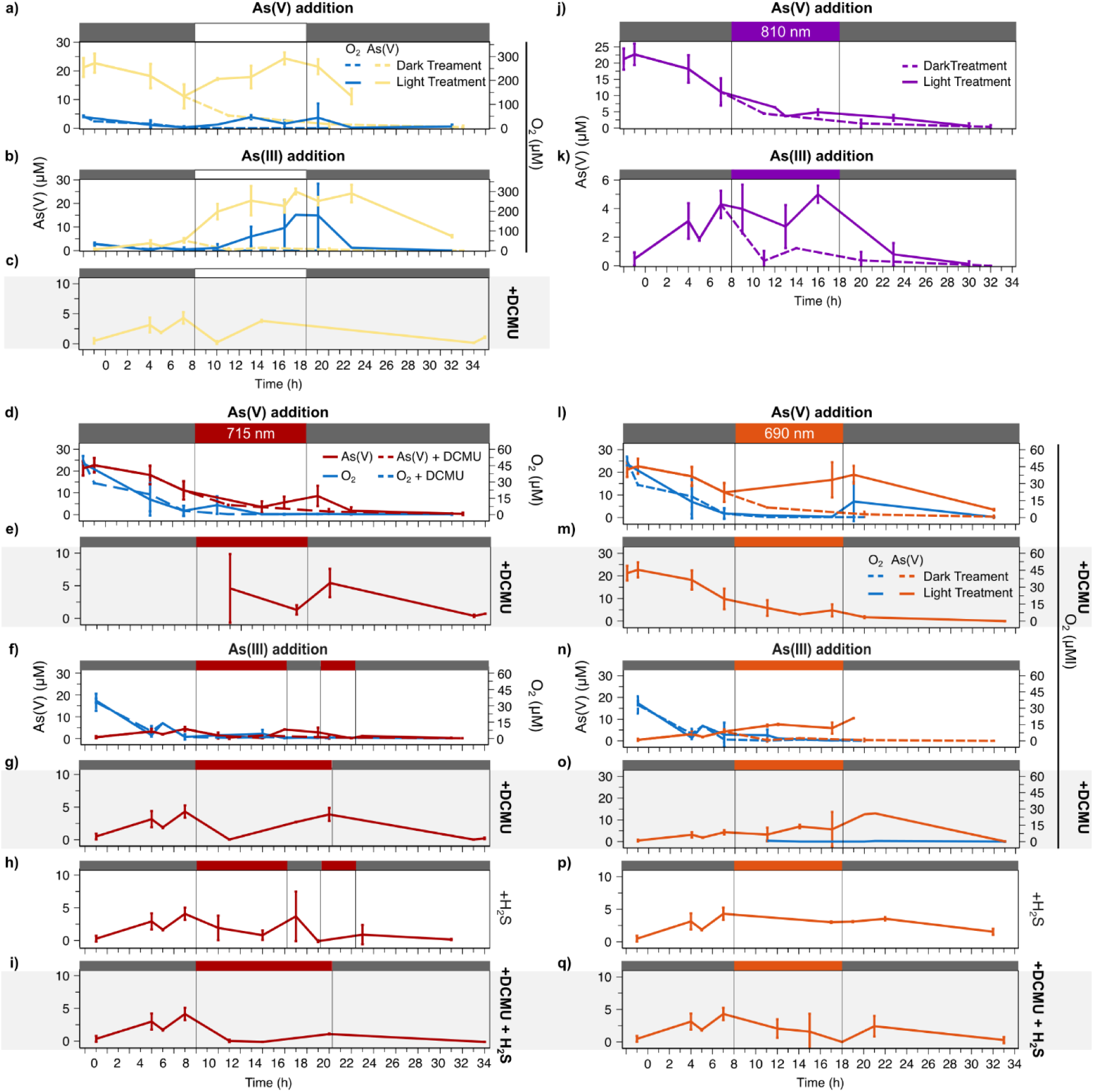
Time series of dissolved As(V) and O_2_ in the overlying water of Pozo Bravo microbial mat incubations, with or without DCMU, an oxygenic photosynthesis inhibitor, treated with white light targetting all phototrophs (a-c), light at 715 nm targetting photosystem I (d-i), light at 810 nm targetting photosystem II (j, k) and light at 690 nm targetting cyanobacterial chl a (l-q). Periods of light are indicated by the colored bars at the top of each graph (white = white light, red = light at 715 nm, purple = light at 810 nm, orange = light at 690 nm). Dark gray bars at the top of the graphs represent periods of darkness. In the As(III) addition experiment at 710 nm light (f and h), incubations without DCMU addition were treated with an extra period of darkness in the afternoon, while in the treatment with DCMU lights were kept on for the entire daylight period. Error bars represent standard deviation, n=3. As(III), As(V), and H_2_S were added to a final concentration of 10 µM.

**Extended data Figure 7:**
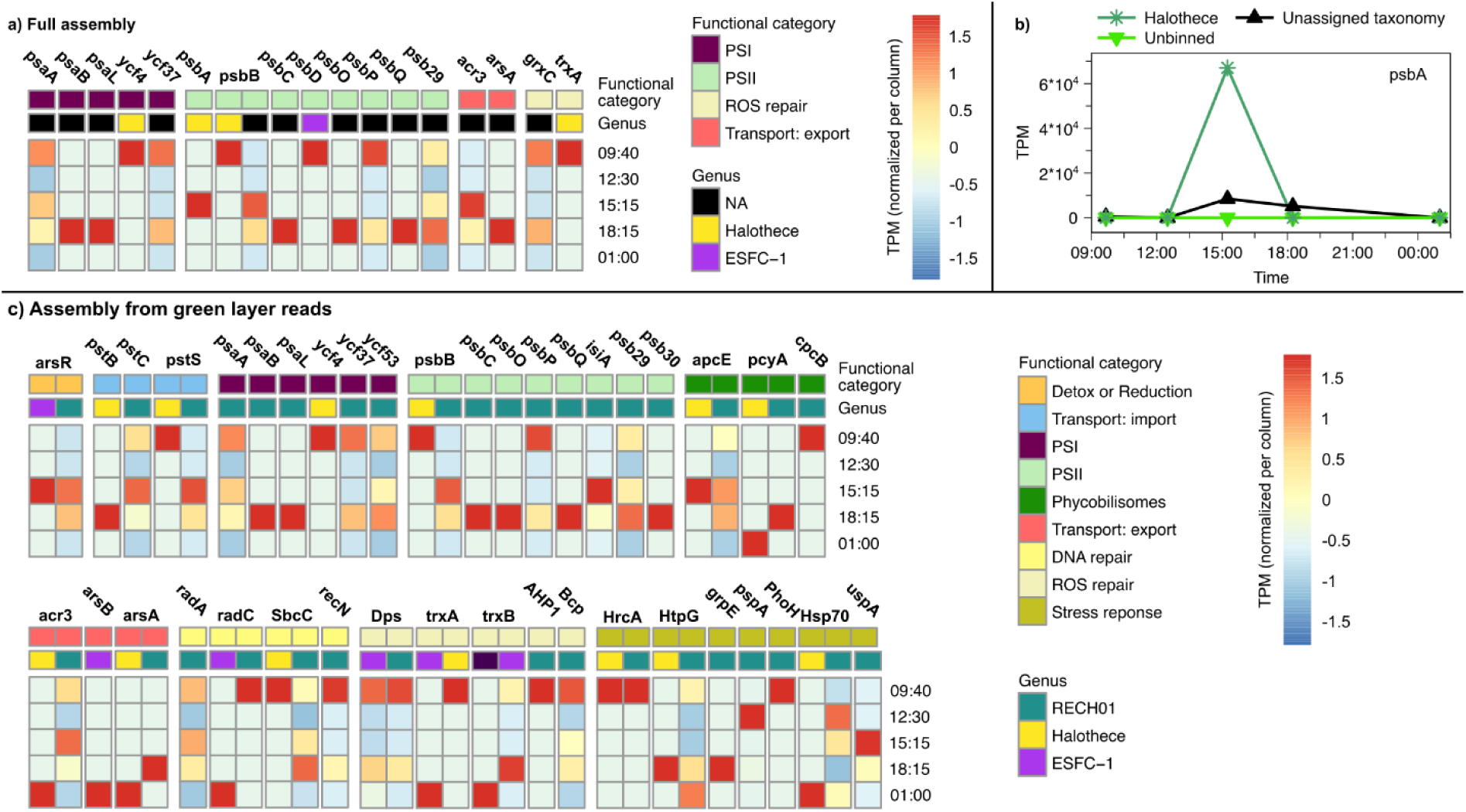
Gene expression, as TPM normalized per column, in bins of order Cyanobacteriia related to arsenic, photosynthesis, DNA repair, and responses to reactive oxygen species (ROS), as well as heat, salinity, and temperature stress responses in a Pozo Bravo microbial mat. In (a) and (c) heatmaps are shown, with higher gene expression in red, lower expression in blue. Normalization was performed due to the varying number of gene copies found per gene. TPM was calculated from metatranscriptomic reads mapped to the metagenome assembly made using only reads from the green microbial mat layer to specifically target cyanobacteria. Two assemblies were made because only a few cyanobacterial genes were found when all samples were used to make the assembly (a), but more cyanobacterial genes could be recovered after the assembly of only the green Pozo Bravo layer (b). In (b) expression of gene psbA, shown as the sum of TPM per genus, is shown. TPM was calculated from metatranscriptomic reads mapped to a metagenomic assembly made from reads of all layers of Pozo Bravo microbial mat. Reads labelled as ‘unassigned taxonomy’ originate from two separate bins to which no taxonomic information could be assigned.

**Extended data Figure 8:**
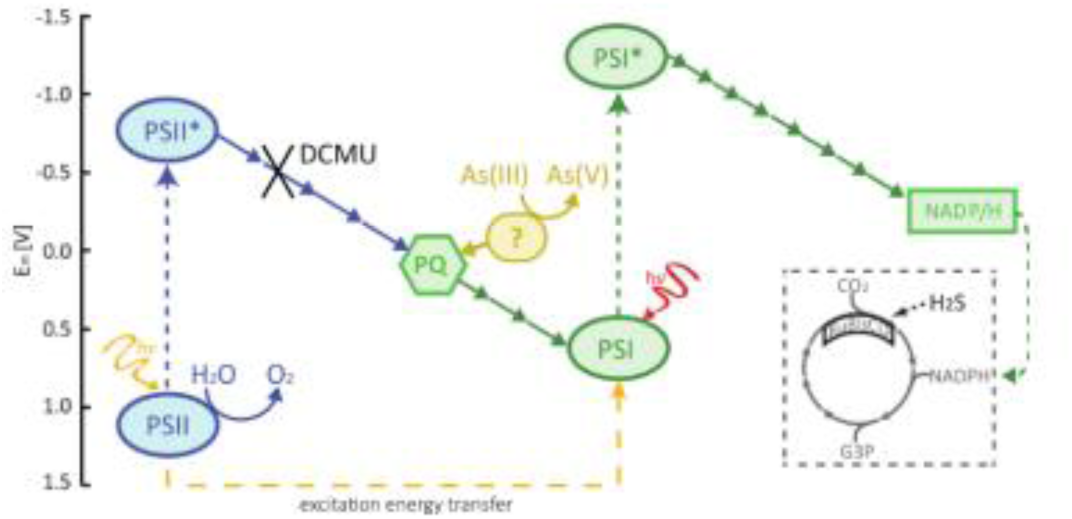
Schematic view of the electron transport during cyanobacterial photosynthesis, indicating where DCMU interrupts electron flow, thus blocking oxygenic photosynthesis. The point of entry for electrons from As(III) must thus be after PSII and only involve PSI (PS= photosystem, PQ= plastoquinone, ?= hypothetical enzyme). Modified from^3^.

