## Supplementary Results for "Cryptic arsenic cycling controls oxygenic photosynthesis in Precambrian-analog microbial mats"

### 1. Microbial mat metagenome analysis

To investigate the vertical distribution of phyla in the mat, we performed metagenomics on slices of a microbial mat. The slices were obtained by separating a piece of mat with a scalpel into the four visible layers (pink, green, red, brown, Fig. 1a), directly after sampling. An assembly was then made using the metagenomic reads from all layers. After assigning the assembled contigs to bins, between 62-69% of the reads from each layer were successfully mapped back to the bins, indicating that the bins give a good overview of the microbial community in each layer (Extended data Fig. 1, Supp. Table S1). Among bins with taxonomic assignments, Pseudomonadota (Alphaproteobacteria and Gammaproteobacteria) was found to be the most abundant phylum across all layers of the microbial mat, followed by Bacteroidota (Bacteroidia and Rhodothermia) and Bacillota (Halanaerobiia, Bacilli, and Clostridia). Chloroflexota (Chloroflexia and Anaerolineae) were also abundant in all layers except for the upper crust, while Deinococcota (Deinococci) were present in a relatively large abundance only in the crust. With the exception of an incomplete Lokiarchaea bin in the red and brown layers, all bins were bacterial. Two cyanobacterial bins, *Halothece sp*. and ESFC-1 (of family *Spirulinaceae*), were recovered from a co-assembly made using all samples. A new assembly was conducted only using reads from the green microbial mat layer to specifically target cyanobacteria. The same cyanobacterial bins, plus RECH01 (of family *Elainellaceae*), were recovered. From the abundance of reads mapped to each bin (Supp. Table S2), ESFC-1 and RECH01 were likely located in the pink and green layers, and Halothece in the green and red layers.

Although pigment absorbance (Fig. 1c) indicated a relatively high abundance of cyanobacteria in the green and red layers of the microbial mat, total cyanobacterial abundance in the mat appears to be underestimated. Low cyanobacterial representation in metagenomes is a well-documented problem[1–4]. This could be due to DNA extraction being hindered by the protective mucilage layer around cyanobacterial cells [1,2]. Low cyanobacterial representation in the metagenome could also be due to the high complexity of the microbial mat community (Fig. 5), and possibly also high strain variation. The relative abundance between the cyanobacterial taxa per mat layer, however, likely reflects the real vertical distribution under the assumption that DNA extraction efficiency is similar for all cyanobacteria through the mat.

***Supplementary Table S1.***

*Genomic bins, filtered at over 55 % completion, selected from a Pozo Bravo metagenomic assembly. The assembly was made using samples from all microbial mat layers ("pink", "green", "red", and "brown").*

| Bin | Completion (%) | Redundancy (%) | Domain | Phylum | Class | Order | Family | Genus | Species |
|---|---|---|---|---|---|---|---|---|---|
| **metabat_bin_31_1_1_3** | 56.34 | 4.23 | NA | NA | NA | NA | NA | NA | NA |
| **metabat_bin_31_1_5_1** | 56.34 | 2.82 | NA | NA | NA | NA | NA | NA | NA |
| **metabat_bin_56_14_10_2** | 56.34 | 1.41 | Bacteria | Proteobacteria | Gammaproteobacteria | Xanthomonadales | Wenzhouxiangellaceae | GCA-2722315 | GCA-2722315 sp002722315 |
| **metabat_bin_56_9** | 56.34 | 5.63 | Bacteria | Bacteroidota | Rhodothermia | Rhodothermales | Salinibacteraceae | Tc-Br11-B2g6-7 | Tc-Br11-B2g6-7 sp001564055 |
| **metabat_bin_31_1_4** | 57.75 | 5.63 | Bacteria | Spirochaetota | Spirochaetia | Spirochaetales | Alkalispirochaetaceae | Alkalispirochaeta | Alkalispirochaeta sp002313505 |
| **metabat_bin_56_14_5_6** | 57.75 | 2.82 | Bacteria | Spirochaetota | Spirochaetia | Spirochaetales | Alkalispirochaetaceae | SLST01 | NA |
| **metabat_bin_56_15_13** | 57.75 | 5.63 | Bacteria | Proteobacteria | Alphaproteobacteria | Caulobacterales | Maricaulaceae | Oceanicaulis | Oceanicaulis sp001657295 |
| **metabat_bin_56_15_6** | 57.75 | 4.23 | NA | NA | NA | NA | NA | NA | NA |
| **metabat_bin_56_5_3** | 57.75 | 1.41 | NA | NA | NA | NA | NA | NA | NA |
| **metabat_bin_79** | 59.15 | 2.82 | Bacteria | Planctomycetota | Phycisphaerae | Phycisphaerales | SM1A02 | GCA-2706885 | GCA-2706885 sp002706885 |
| **metabat_bin_132** | 60.56 | 2.82 | NA | NA | NA | NA | NA | NA | NA |
| **metabat_bin_31_5_4___1** | 60.56 | 2.82 | NA | NA | NA | NA | NA | NA | NA |
| **metabat_bin_122** | 61.97 | 2.82 | NA | NA | NA | NA | NA | NA | NA |
| **metabat_bin_51** | 61.97 | 1.41 | NA | NA | NA | NA | NA | NA | NA |
| **metabat_bin_105** | 63.38 | 2.82 | Bacteria | Firmicutes | Bacilli | Bacillales | Halobacillaceae | Halobacillus | NA |
| **metabat_bin_56_14_9_1** | 63.38 | 1.41 | Bacteria | Proteobacteria | Gammaproteobacteria | Chromatiales | Chromatiaceae | PWTO01 | PWTO01 sp003562815 |
| **metabat_bin_56_5_2** | 63.38 | 2.82 | Bacteria | Proteobacteria | Gammaproteobacteria | Chromatiales | Chromatiaceae | PWTO01 | PWTO01 sp003562815 |
| **metabat_bin_1** | 64.79 | 7.04 | Bacteria | Firmicutes | Halanaerobiia | Halanaerobiales | Halanaerobiaceae | Halanaerobium | NA |
| **metabat_bin_131** | 64.79 | 0.00 | NA | NA | NA | NA | NA | NA | NA |
| **metabat_bin_171_1** | 64.79 | 0.00 | NA | NA | NA | NA | NA | NA | NA |
| **metabat_bin_31_5_5__2** | 64.79 | 0.00 | Bacteria | Chloroflexota | Anaerolineae | Anaerolineales | Anaerolineaceae | Brevefilum | Brevefilum sp007119455 |
| **metabat_bin_108** | 66.20 | 2.82 | Bacteria | Proteobacteria | Alphaproteobacteria | Rhodobacterales | Rhodobacteraceae | HLUCCA09 | HLUCCA09 sp003340565 |
| **metabat_bin_31_4_2_1** | 67.61 | 4.23 | NA | NA | NA | NA | NA | NA | NA |
| **metabat_bin_120** | 69.01 | 7.04 | Bacteria | Proteobacteria | Alphaproteobacteria | Rhodobacterales | Rhodobacteraceae | NA | NA |
| **metabat_bin_126** | 69.01 | 4.23 | Bacteria | Proteobacteria | Alphaproteobacteria | Rhodobacterales | Rhodobacteraceae | Rubrimonas | Rubrimonas cliftonensis |
| **metabat_bin_31_5_1_4** | 69.01 | 2.82 | Bacteria | Chloroflexota | Anaerolineae | SBR1031 | A4b | J038 | J038 sp003696565 |

| Bin | Completion (%) | Redundancy (%) | Domain | Phylum | Class | Order | Family | Genus | Species |
|---|---|---|---|---|---|---|---|---|---|
| **metabat_bin_15** | 70.42 | 1.41 | NA | NA | NA | NA | NA | NA | NA |
| **metabat_bin_56_14_12_1** | 71.83 | 5.63 | NA | NA | NA | NA | NA | NA | NA |
| **metabat_bin_27** | 73.24 | 0.00 | Bacteria | Proteobacteria | Alphaproteobacteria | Rhodospirillales | Rhodospirillaceae | Rhodospira | Rhodospira trueperi |
| **metabat_bin_31_5_1_2** | 73.24 | 4.23 | Bacteria | Proteobacteria | Gammaproteobacteria | Pseudomonadales | Oleiphilaceae | Marinobacter | NA |
| **metabat_bin_56_11_1** | 73.24 | 2.82 | Bacteria | Chloroflexota | Chloroflexia | Chloroflexales | Chloroflexaceae | NA | NA |
| **metabat_bin_94** | 74.65 | 0.00 | Bacteria | Proteobacteria | Alphaproteobacteria | Rhodobacterales | Rhodobacteraceae | Rubrimonas | Rubrimonas cliftonensis |
| **metabat_bin_145** | 76.06 | 0.00 | Bacteria | Proteobacteria | Alphaproteobacteria | Kiloniellales | Rhodovibrionaceae | Rhodovibrio | Rhodovibrio salinarum |
| **metabat_bin_56_14_13_1** | 76.06 | 5.63 | Bacteria | Chloroflexota | Anaerolineae | SBR1031 | A4b | OLB15 | OLB15 sp001567085 |
| **metabat_bin_56_15_5** | 76.06 | 2.82 | Bacteria | Proteobacteria | Alphaproteobacteria | Rhodobacterales | Rhodobacteraceae | Roseovarius | Roseovarius sp007125415 |
| **metabat_bin_56_5_1** | 76.06 | 1.41 | NA | NA | NA | NA | NA | NA | NA |
| **metabat_bin_124** | 77.46 | 4.23 | NA | NA | NA | NA | NA | NA | NA |
| **metabat_bin_31_5_5__1** | 77.46 | 1.41 | Bacteria | Chloroflexota | Anaerolineae | Anaerolineales | Anaerolineaceae | Brevefilum | Brevefilum sp007119455 |
| **metabat_bin_37** | 77.63 | 5.26 | NA | NA | NA | NA | NA | NA | NA |
| **metabat_bin_169** | 78.87 | 2.82 | NA | NA | NA | NA | NA | NA | NA |
| **metabat_bin_56_14_2** | 80.28 | 2.82 | NA | NA | NA | NA | NA | NA | NA |
| **metabat_bin_31_4_1** | 81.69 | 2.82 | Bacteria | Deinococcota | Deinococci | Deinococcales | Trueperaceae | NA | NA |
| **metabat_bin_118** | 83.10 | 1.41 | NA | NA | NA | NA | NA | NA | NA |
| **metabat_bin_56_3** | 83.10 | 2.82 | Bacteria | Bacteroidota | Rhodothermia | Rhodothermales | Salinibacteraceae | Longibacter | Longibacter salinarum |
| **metabat_bin_38** | 84.51 | 1.41 | NA | NA | NA | NA | NA | NA | NA |
| **metabat_bin_56_14_1** | 84.51 | 5.63 | Bacteria | Spirochaetota | Spirochaetia | Spirochaetales | Alkalispirochaetaceae | SLST01 | SLST01 sp007127335 |
| **metabat_bin_31_5_1_3** | 85.92 | 4.23 | NA | NA | NA | NA | NA | NA | NA |
| **metabat_bin_56_14_5_2** | 85.92 | 4.23 | Bacteria | Chloroflexota | Chloroflexia | Chloroflexales | Chloroflexaceae | Viridilinea | Viridilinea mediisalina |
| **metabat_bin_56_6** | 85.92 | 7.04 | Bacteria | Spirochaetota | Spirochaetia | Spirochaetales | Alkalispirochaetaceae | SLST01 | NA |
| **metabat_bin_144** | 87.32 | 5.63 | Bacteria | Proteobacteria | Alphaproteobacteria | Rhodobacterales | Rhodobacteraceae | NA | NA |
| **metabat_bin_56_12** | 87.32 | 0.00 | NA | NA | NA | NA | NA | NA | NA |
| **metabat_bin_77** | 87.32 | 4.23 | NA | NA | NA | NA | NA | NA | NA |
| **metabat_bin_156** | 88.73 | 1.41 | NA | NA | NA | NA | NA | NA | NA |
| **metabat_bin_56_14_8** | 88.73 | 5.63 | Bacteria | Proteobacteria | Gammaproteobacteria | NA | NA | NA | NA |
| **metabat_bin_56_15_1** | 88.73 | 0.00 | Bacteria | Deinococcota | Deinococci | Deinococcales | Trueperaceae | NA | NA |
| **metabat_bin_56_15_2** | 88.73 | 2.82 | Bacteria | Proteobacteria | Alphaproteobacteria | Rhizobiales | Beijerinckiaceae | Salinarimonas | Salinarimonas sp900094735 |
| **metabat_bin_56_4_1** | 88.73 | 0.00 | Bacteria | Proteobacteria | Alphaproteobacteria | Rhizobiales | Rhodomicrobiaceae | Dichotomicrobium | Dichotomicrobium thermohalophilum |

| Bin | Completion (%) | Redundancy (%) | Domain | Phylum | Class | Order | Family | Genus | Species |
|---|---|---|---|---|---|---|---|---|---|
| **metabat_bin_56_10** | 90.14 | 4.23 | Bacteria | Proteobacteria | Gammaproteobacteria | Chromatiales | Chromatiaceae | PWTO01 | PWTO01 sp003562815 |
| **metabat_bin_56_14_5_3** | 90.14 | 2.82 | Bacteria | Proteobacteria | Gammaproteobacteria | Xanthomonadales | Wenzhouxiangellaceae | Wenzhouxiangella | Wenzhouxiangella marina |
| **metabat_bin_66** | 90.14 | 0.00 | Bacteria | Proteobacteria | Gammaproteobacteria | Legionellales | Legionellaceae | Legionella | Legionella geestiana |
| **metabat_bin_17** | 91.55 | 5.63 | Bacteria | Cyanobacteria | Cyanobacteriia | Cyanobacteriales | Rubidibacteraceae | Halothece | Halothece sp000317635 |
| **metabat_bin_173_1** | 91.55 | 4.23 | Bacteria | Spirochaetota | Spirochaetia | Spirochaetales | Alkalispirochaetaceae | NA | NA |
| **metabat_bin_75** | 91.55 | 2.82 | Bacteria | Bacteroidota | Bacteroidia | Bacteroidales | UBA12077 | UBA12077 | UBA12077 sp002869305 |
| **metabat_bin_146** | 92.96 | 1.41 | NA | NA | NA | NA | NA | NA | NA |
| **metabat_bin_56_2** | 92.96 | 8.45 | Bacteria | NA | NA | NA | NA | NA | NA |
| **metabat_bin_136** | 94.37 | 0.00 | Bacteria | Bacteroidota | Bacteroidia | Flavobacteriales | T3Sed10-241 | T3Sed10-241 | NA |
| **metabat_bin_168** | 94.37 | 2.82 | NA | NA | NA | NA | NA | NA | NA |
| **metabat_bin_171_2** | 94.37 | 2.82 | NA | NA | NA | NA | NA | NA | NA |
| **metabat_bin_31_1_2** | 94.37 | 4.23 | NA | NA | NA | NA | NA | NA | NA |
| **metabat_bin_35** | 94.37 | 8.45 | NA | NA | NA | NA | NA | NA | NA |
| **metabat_bin_56_13** | 94.37 | 7.04 | NA | NA | NA | NA | NA | NA | NA |
| **metabat_bin_56_7** | 94.37 | 1.41 | Bacteria | Bacteroidota | Rhodothermia | Rhodothermales | Salinibacteraceae | Longibacter | Longibacter salinarum |
| **metabat_bin_20** | 95.77 | 0.00 | Bacteria | Bacteroidota | Bacteroidia | Bacteroidales | NA | NA | NA |
| **metabat_bin_31_1_1** | 95.77 | 4.23 | Bacteria | Cyanobacteria | Cyanobacteriia | NA | NA | NA | NA |
| **metabat_bin_165** | 97.18 | 4.23 | NA | NA | NA | NA | NA | NA | NA |
| **metabat_bin_86** | 97.18 | 8.45 | NA | NA | NA | NA | NA | NA | NA |
| **metabat_bin_87** | 97.18 | 5.63 | NA | NA | NA | NA | NA | NA | NA |
| **metabat_bin_138** | 98.59 | 1.41 | Bacteria | Bacteroidota | Bacteroidia | Bacteroidales | NA | NA | NA |
| **metabat_bin_14** | 98.59 | 1.41 | NA | NA | NA | NA | NA | NA | NA |
| **metabat_bin_149** | 98.59 | 1.41 | Bacteria | Bacteroidota | Bacteroidia | Cytophagales | Cyclobacteriaceae | NA | NA |
| **metabat_bin_52** | 98.59 | 1.41 | NA | NA | NA | NA | NA | NA | NA |
| **metabat_bin_54** | 98.59 | 5.63 | Bacteria | Bacteroidota | Bacteroidia | Bacteroidales | Salinivirgaceae | Salinivirga | Salinivirga cyanobacteriivorans |
| **metabat_bin_56_8** | 98.59 | 1.41 | Bacteria | Bacteroidota | Rhodothermia | Rhodothermales | Salinibacteraceae | Longibacter | Longibacter salinarum |
| **metabat_bin_7** | 98.59 | 1.41 | Bacteria | Firmicutes | Bacilli | Bacillales | Amphibacillaceae | Halolactibacillus | NA |
| **metabat_bin_92** | 98.59 | 4.23 | Bacteria | Bacteroidota | Rhodothermia | Rhodothermales | Salinibacteraceae | NA | NA |
| **metabat_bin_106** | 100.00 | 4.23 | NA | NA | NA | NA | NA | NA | NA |
| **metabat_bin_50** | 100.00 | 4.23 | Bacteria | Firmicutes | Halanaerobiia | Halobacteroidales | Halobacteroidaceae | Orenia | Orenia marismortui |

***Supplementary Table S2.***

*Total abundance (%) of cyanobacterial metagenomic reads that were mapped back onto the metagenomic reads of each microbial mat layer ("pink", "green", "red", and "brown"). Based on microsensor measurements of oxygen production, cyanobacteria were expected in the green and red layers. The cyanobacterial reads were obtained from the metagenomic assembly made using only reads from the green layer, because cyanobacterial reads were difficult to recover from the assembly made using all samples ("pink", "green", "red", and "brown").*

| Cyanobacterial genus | Pink | Green | Red | Brown |
|---|---|---|---|---|
| ESFC-1 | 0.34 | 0.27 | 0.01 | 0.00 |
| Halothece | 0.25 | 1.05 | 0.54 | 0.03 |
| RECH01 | 0.40 | 0.52 | 0.07 | 0.09 |

### 2. Arsenic redox pathways in the metagenome

We used our metagenome data to screen for genes involved in arsenic metabolism. A great variety of genes directly related to arsenic metabolism are present in the Pozo Bravo microbial mat (Fig. 1). The roles of these genes are discussed in more detail below and shown in Fig. 1, and Supp. Table S3.

Several distinct routes for microbial arsenic detoxification exist, broadly divided into three steps. Initially, arsenic gets imported into microbial cells via specific transporters, then it is chemically modified by specific detoxification systems, and then finally exported using dedicated efflux systems. First, we focused on arsenic import pathways. As(V) and As(III) can enter the cell through different mechanisms. As(III) (in the protonated form $As(OH)_3$) can enter the cell by diffusion across the membrane or through aquaglyceroporin channels (*aqps*), the genes for which were found in the metagenome [5,6]. As(V) can be imported through phosphate channels [7]. Almost all components (*pstSCAB*) of the high-affinity phosphate import channel were found. The pst system is also highly selective for phosphate (up to $10^3$), which results from crucial bonds in the transporter being affected by the slightly larger size of As(V) [8]. Thus, the use of pst indicates an adaptation to limit As(V) intake (see Supp. Results 5) [7–9].

Once inside the cell, arsenic can be metabolized for energy generation or detoxified through a series of redox reactions. The genes found in the Pozo Bravo metagenome were summarized into three putative detoxification-related pathways (Fig. 3): The first detoxification pathway is based on an As(V)-transferring gap3/gadph, and arsJ, which were both found in the metagenome. The second pathway involves reduction of As(V) to As(III) via arsC1, arsC2 and mrx1. The genes encoding for arsC1 and arsC2 were not recovered from the metagenome. Yet, their presence is required for functionality of mrx1 (Fig. 3, for the full Mrx1 cycle see Ordóñez et al. 2009 [10]. While this pathway has thus far exclusively been assigned a detoxification function, the third pathway involving As(V) reduction mediated by the product of the genes *arsRCDAB* and *acr3*, has been hypothesized to be used in As(V) respiration [11]. In all other studies of genomes and metagenomes in the Puna region, an environment similar to Pozo Bravo, *acr3* was the predominant As(V) reduction gene [12–14]. No other respiratory *arrA* or *arrB* As(V) reductases were found. As(III) oxidation is mediated by the products of *aioA* and *aioB*, coupled to the reduction of either cytochrome c or azurin, as part of processes such as AP [15]. Both *aioA* and *aioB* were found in reads from all layers of the metagenome. Other arsenite oxidation genes (*arxA*) were not found in the metagenome. Overall, the metagenomic analysis revealed a significant abundance of arsenic-related genes in all layers of the Pozo Bravo microbial mat, thus enabling bacteria to thrive under these harsh conditions.

***Supplementary Table S3.***

*Summary of the genes listed in the main text, based on previously reported functions*[10,16,25–29,17–24].

| Gene | Relevance | Functional category | Synonyms (literature) |
|---|---|---|---|
| **acr3** | As(III) exporter | Transport: export | |
| **AHP1** | peroxiredoxin-5 | ROS repair | |
| **AhpC** | part of alkyl hydroperoxide reductase | ROS repair | |
| **AhpF** | part of alkyl hydroperoxide reductase | ROS repair | |
| **aioA** | part of As(III) oxidase AioAB | Oxidation | aoxA, aroB |
| **aioB** | part of As(III) oxidase AioAB | Oxidation | aoxB, aroA |
| **apcE** | part of phycobilisome | Phycobilisomes | |
| **aqps** | As(III) import channel | Transport: import | |
| **arrA** | respiratory As(V) reductase | Reduction | |
| **arrB** | respiratory As(V) reductase | Reduction | |
| **arsA** | part of the transporter ArsAB, which exports As(II) | Transport: export | |
| **arsB** | part of the transporter ArsAB, which exports As(II). Can function without ArsA component | Transport: export | |
| **arsC** | As(V) reductase | Reduction | |
| **arsC1, arsC2** | As(V) mycothioltransferase, which yields As(V)-mycothiol | Detoxification | |
| **arsD** | As(V) chaperone to ArsA component of ArsAB transporter | Detoxification | |
| **arsJ** | 1-arsono-3-phospho-D-glycerate exporter | Transport: export | |
| **arsR** | inhibits *ars* genes under low [As(III)] | Regulation of detox/reduction | |
| **arxA** | part of As(III) oxidase ArxAB | Oxidation | |
| **arxB** | part of As(III) oxidase ArxAB | Oxidation | |

| Gene | Relevance | Functional category | Synonyms (literature) |
|---|---|---|---|
| **Bcp** | thiol peroxidase | ROS repair | |
| **cpcB** | part of phycobilisome | Phycobilisomes | |
| **Dps** | ferritin-like DNA-binding proteins from starved cells | ROS repair | |
| **gapdh** | If As(V)-transferring, catalyzes As(V)->1-arsono-3-phospho-D-glycerate | Detoxification | gap3 |
| **grpE** | involved in heat shock response with Hsp70 | Stress reponse | |
| **grxC** | As(V)-related glutaredoxin, reduces As(V). May also be involved in ROS | Reduction / ROS repair | |
| **grxD** | Glutaredoxin | ROS repair | |
| **HrcA** | heat shock protein | Stress reponse | |
| **Hsp70** | heat shock protein | Stress reponse | DnaK |
| **htpG** | heat shock protein | Stress reponse | |
| **isiA** | encodes D2 protein in Photosystem II, under Fe-limiting conditions | Photosystem II | |
| **mrx1** | Removes mycothiol component of As(V)-mycothiol, yields As(III) for export | Reduction | |
| **nrdH** | oxidoreductases similar to glutaredoxins, but function like thioredoxins | ROS repair | |
| **pcyA** | part of phycobilisome | Phycobilisomes | |
| **PhoH** | makes ATPase under PO4 starvation | Stress reponse | |
| **pitA, pitB** | As(V) importer- lower PO4 affinity than pit | Transport: import | |
| **psbA** | encodes D1 protein in Photosystem II, D1 type adapted to environmental conditions | Photosystem II | |
| **psbA, psaB, psaL** | part of Photosystem I | Photosystem I | |
| **psbB, psbC, pdbD, psbO, psbP, psbQ, psb29, psb30** | part of Photosystem II | Photosystem II | |
| **pspA** | lipid membrane repair | Stress reponse | |
| **pstS, pstC, pstA, pstB, pho U** | As(V) importer- higher phosphate affinity than pit | Transport: import | |
| **pufL** | part of Reaction Center II | Photosynthetic Reaction Center II | |

| Gene | Relevance | Functional category | Synonyms (literature) |
|---|---|---|---|
| **pufM** | part of Reaction Center II | Photosynthetic Reaction Center II | |
| **radA** | linked to repair of UV damage | DNA repair | |
| **radC** | linked to repair of UV damage | DNA repair | |
| **recN** | general DNA repair | DNA repair | |
| **SbcC** | general DNA repair | DNA repair | |
| **Tpx** | thiol peroxidase | ROS repair | Prx |
| **trx** | thioredoxin | ROS repair | |
| **trxA** | thioredoxin | ROS repair | |
| **trxA** | thioredoxin | ROS repair | |
| **trxB** | thioredoxin reductase | ROS repair | |
| **uspA** | universal strass protein | Stress reponse | |
| **ycf4, ycf37, ycf 53** | part of Photosystem I | Photosystem I | |

### 3. Diel dynamics of Fe and $H_2S$

Analysis of porewater collected over a diel cycle directly in the field, showed no significant dynamics of iron concentration and speciation, due to the low iron concentration (< 1 µM) and high heterogeneity between replicates (data not shown). $H_2S$ concentration was low (~20 µM at 1 cm depth based on ex-situ measurements, Supplementary Fig. S1). Unfortunately, in-situ $H_2S$ depth profiling was only successful for short intermittent phases of the day in one spot and difficult to interpret due to strong signal drifts after calibration. Measurement of pH by microsensors failed entirely due to the hard texture of the mat, which impeded calculation of total sulfide. Yet, the depth of the $H_2S$ consumption zone was light-dependent (Supplementary Fig S1). $H_2S$ was predominantly depleted at 3-4 mm depth during the day, but reached the uppermost layer, the oxic interface, in the evening. Due to the absence of $O_2$ below 3 mm in the afternoon, $H_2S$ oxidation with photosynthetically produced $O_2$ is unlikely.

To further differentiate between $H_2S$-driven photosynthesis or an effect of pH shifts on $H_2S$ concentration, we performed ex-situ measurements and assessed light-dependent dynamics of $H_2S$ concentration at each depth, which revealed anoxygenic photosynthetic activity at ~4 mm depth (Fig S1). However, rates were very low, suggesting that the light-dependent $H_2S$ dynamics in situ were largely due to pH changes driven by a different process.

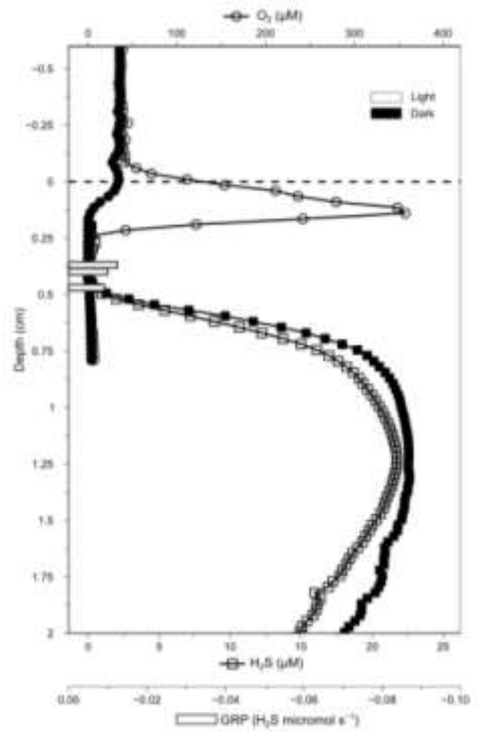

***Supplementary Figure S1.***

*Microsensor profiles of $O_2$ and $H_2S$ concentration from a Pozo Bravo microbial mat, taken ex-situ. Open symbols indicate measurements taken under light, closed symbols indicate measurements taken during darkness. Bars indicate the Gross Rate of anoxygenic ($H_2S$-driven) Photosynthesis (GRP).*

### 4. Metagenomic evidence for concurrent arsenic oxidation and reduction

We used metatranscriptomes from Pozo Bravo to determine diel dynamics in transcription of genes related to arsenic metabolism. Specifically, we investigated photosynthetic gene transcription in the afternoon, and whether transcription of genes related to arsenite oxidation increased during the same period. For this purpose, we mapped transcriptomic reads from in-situ mat samples obtained over a diel cycle to the metagenomic bins (Fig. 5).

The highest read numbers mapped to arsenic genes were present in Alpha- and Gamma-proteobacterial bins (Fig. 5), which stimulated in-depth analysis of diel transcription by these groups. We found that As(III) oxidation was likely mainly driven by *aioAB* transcribed by taxa among the Rhodobacterales and Chromatiales, prominent representatives of purple photosynthetic bacteria. Specifically, *aioA* encoding for the large subunit of arsenite oxidase, was transcribed at all timepoints in Rhodobacterales (Alphaproteobacteria) (Fig. 5). The highest *aioA* transcription, and thus potentially As(III) oxidation, happened during the day (9 am - 6 pm) although most of the arsenic present in porewater was As(V) (Fig. 5). Transcription of *aioB*, which encodes for the small subunit of the same As(III) oxidase, was also found during 9 am to 3 pm, in line with high *aioAB* activity during this portion of the day. During daylight hours, Rhodobacterales expressing *aioAB* also transcribed *pufL* and *pufM* (Supp. Fig. S2), which encode for components of a type II photosynthetic reaction center, thus supporting arsenite-driven anoxygenic photosynthesis. Similar patterns of *aioAB* and *pufLM* transcription were observed in Rhizobiales, mainly during the morning and late afternoon (Supp. Fig. S2, Fig. 5b).

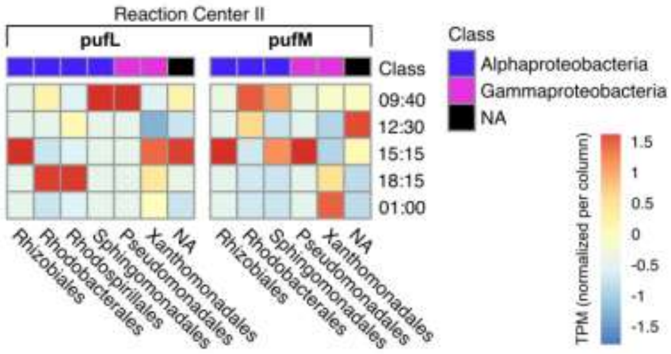

***Supplementary Figure S2***. *Gene expression in TPM normalized per bacterial class and gene (i.e.- normalized per column), of the pufL and pufM components of photosynthetic reaction center II.*

Gammaproteobacteria were separated into three main orders: Chromatiales, Pseudomonadales and Xanthomonadales (Fig. 5b). Transcription of *pufLM* was found in both Pseudomonadales and Xanthomonadales during the day as well as the night. However, unlike in Alphaproteobacteria, transcription of *aioA* primarily took place during the night, except for Chromatiales that showed transcription at 3 pm. Transcription of *aioB* was not found. This could suggest non-photosynthetic As(III) oxidation at night, i.e.- coupled to nitrate reduction [30,31], in analogy to chemolithoautotrophic sulfide oxidation by purple sulfur bacteria commonly observed in other mat ecosystems [32].

The potential As(III)-oxidizing purple anoxygenic phototrophs among the Alphaproteobacteria appeared to transition to As(V) reduction at low light and in the night. Namely, in Rhodobacterales, the transcription of 1-arseno-3-phosphoglycerate via As(V)-transferring GAPDH and *arsJ* peaked at night, and As(V) reduction and As(III) excretion via *arsC* and *arsB*, occurred in the morning. Most Gammaproteobacteria did not show such pattern. Instead, *arsC*, *acr3* and *arsB* were continuously expressed. Chromatiales, however, co-expressed *aioA* and genes involved in excretion of intracellular As(III) (*arsA*, *arsB*) during the day. This general pattern of alternating key players in As(V) reduction held true across the complete metatranscriptome. Most prominently, the relative contribution of bacterial classes driving arsenate reduction via *arsC* changed per timepoint (Fig. 5a). Transcription of *arsC* by Gammaproteobacteria was high and relatively constant, while *arsC* transcription at 9 am was highest by Alphaproteobacteria, and at noon by Bacteroidota and Chloroflexota. Spirochaetia was the only known bacterial class to express *arsC* at night (1 am), but some transcription was also found in unidentified bins. Transcription in unbinned reads was negligible. While the key players may alternate over a diel cycle, intriguingly, the general *arsC* transcription was highest during daylight hours (Fig. 5a), consistent with pronounced cryptic redox cycling of arsenic.
